# Irc3 of the thermotolerant yeast *Ogataea polymorpha* is a branched-DNA-specific mitochondrial helicase

**DOI:** 10.1101/2022.03.21.485104

**Authors:** Vlad-Julian Piljukov, Sirelin Sillamaa, Tiina Sedman, Natalja Garber, Margus Rätsep, Arvi Freiberg, Juhan Sedman

## Abstract

The *Saccharomyces cerevisiae* Irc3 protein is a mitochondrial Superfamily II DNA helicase that, according to genomic data, is conserved in different yeast species. Here we characterize Irc3 helicase from the thermotolerant yeast *Ogataea polymorpha* (Irc3_op_) that throughout its helicase motor domain has approximately 82% similarity with the *S. cerevisiae* protein. Irc3_op_ retains enzymatic activity at considerably higher temperatures than Irc3_sc_, displaying the fastest rate of ATP hydrolysis at 41 °C and a Tm of 45.3 °C for its secondary structure melting. We demonstrate that Irc3_op_ is a structure-specific DNA helicase translocating on both single- and double-stranded DNA molecules. Like the homolog of *S. cerevisiae*, Irc3_op_ can unwind only DNA molecules that contain branched structures. Different DNA molecules containing three- and four-way branches are utilized by Irc3_op_ as unwinding substrates. Importantly, the preferred unwinding substrate of Irc3_op_ is a DNA fork containing the nascent lagging strand, suggesting a possible role in replication restart following a block in leading strand polymerization. A lower unwinding efficiency of four-way branched DNA molecules could explain why Irc3_op_ only partially complements the *irc3*Δ phenotype in *S. cerevisiae*.

## Introduction

Helicases, mechanochemical enzymes that unwind or remodel nucleic acid molecules in an ATP-dependent manner are involved in all aspects of nucleic acid metabolism, including DNA replication, recombination and repair [1,2]. Mutations in several helicases are related to human disease, causing premature ageing syndromes, most notable being Bloom’s and Werner syndromes [3,4], or increasing the risk of developing cancer [5–7]. In eukaryotic cells, where the majority of DNA helicases are involved in nuclear DNA metabolism, a small number of helicases are targeted to mitochondria, semiautonomous organelles with their own genomic DNA. The mitochondrial genome encodes for 8-13 essential peptides of respiratory chain complexes and thus, the maintenance of mitochondrial genome stability is crucial for cellular respiration. The proteins involved in mitochondrial genome stability, including mitochondrial helicases, are encoded by the nuclear genome, synthesized by cytosolic ribosomes and imported into mitochondria by a specialized transport machinery [8].

Curiously, eukaryotic organisms belonging to different phylogenetic lineages appear to rely on distinct strategies to ensure the stability of their mitochondrial genome and utilize different sets of protein factors to replicate the mitochondrial genome. Through the analysis of mutant yeast strains that display defective mitochondrial DNA metabolism, three DNA helicases (Pif1, Hmi1 and Irc3) have been identified in *Saccharomyces cerevisiae* [9–11]. Hmi1 and Irc3 are targeted exclusively to mitochondria and have no homologs in human cells. Pif1 helicase is conserved from yeast to human and has two isoforms, one targeted to the mitochondria and the other to the nucleus [12,13]. Conversely, the hexameric DNA helicase Twinkle, which is required for the stability of human mtDNA, is not found in yeast mitochondria [14]. DNA helicases iunique to yeast mitochondria make them potential targets of novel antifungal drugs.

Irc3 of the budding yeast *S. cerevisiae* (Irc3_sc_) was first described by our lab as a dsDNA-stimulated ATPase involved in mitochondrial DNA metabolism [11]. Irc3_sc_ belongs to the Superfamily II of helicases and is required for the stability of the *wt* mitochondrial genome, as the disruption of Irc3_sc_ leads to a rapid decline in respiratory activity and to the loss of the *wt* mitochondrial genome in the yeast [11]. The ATP hydrolysis activity of *S. cerevisiae* Irc3_sc_ is stimulated, in addition to dsDNA, also by branched DNA molecules [15]. However, for single-stranded DNA or RNA cofactors, virtually no stimulation of ATP hydrolysis activity can be detected [15]. Irc3_sc_ helicase efficiently unwinds branched DNA substrates, including four-way structures mimicking a Holliday junction and the nascent strands of three-way structures mimicking stalled replication forks [15,16]. In contrast, no DNA unwinding activity of Irc3 can be detected with linear dsDNA molecules [15]. Disruption of Irc3_sc_ in *S. cerevisiae* can partially be complemented by the bacterial RecG helicase targeted to mitochondria; this and the enzymatic properties of Irc3_sc_ support the idea that Irc3 and RecG might fulfill similar biochemical functions [15]. Irc3_sc_ is rapidly inactivated at temperatures exceeding 30 °C and the limited thermal stability has impeded biochemical and structural studies of the protein. We therefore initiated a search of Irc3_sc_ homologs in thermotolerant yeast species. Here we demonstrate that Irc3_op_, encoded by the genome of *Ogataea polymorpha*, is a branched-DNA-specific helicase that remains fully active above 40°C. While the enzymatic properties of the two proteins are similar, the preferred unwinding substrate of Irc3_op_ appears to be the nascent lagging strand at the replication fork whereas Irc3_sc_ is most active on substrates mimicking Holliday junctions with a mobile core structure.

## Experimental procedures

### DNA

The oligonucleotides used in this study were purchased from TAG Copenhagen A/S and are listed in Supplementary Table S1. The DNA cofactors used in the assays of ATP hydrolysis were prepared by annealing complementary oligonucleotides as previously described [15].^32^P-labelled DNA substrates for unwinding assays were prepared by labeling specific oligonucleotides at the 5’-end with polynucleotide kinase and γ-^32^P-ATP as previously described [15]. DNA cofactors used in fluorescence anisotropy measurement experiments were prepared by annealing complementary oligonucleotides, one of which contained a fluorescein moiety at the 5’-terminus (TAG Copenhagen A/S).

The plasmids used for complementation studies of Irc3 deletion in *S. cerevisiae* were based on pRS315 as described in Sedman et al., 2016. pRS315-Irc3_op_ was prepared by replacing the IRC3 open reading frame of *S. cerevisiae* with the coding sequence of the IRC3 homolog from *O. polymorpha NCYC 495 leu1.1* (e_gw1.4.413.1, MycoCosm database [17]). The bacterial expression construct of Irc3_op_ was based on pET24d-His10-SUMO [18] and encoded the amino acid residues 20-604 of the *O. polymorpha* Irc3 preprotein, corresponding to the predicted mature 68.6 kDa protein that lacks the cleavable mitochondrial targeting peptide.

The single-stranded form of the phagemid pGEM-11Zf+ and the superspiral pGEM-11Zf+ (Promega, Madison, WI, USA) were used as cofactors in ATP hydrolysis activity assays.

### Proteins

Irc3 of *O. polymorpha* was overexpressed as a His10-SUMO-Irc3 fusion protein in *E. coli* BL21 DE3 (RIL) transformed with the expression plasmid. Bacteria were grown to OD_600_ ≈ 0.8 in LB media and induced with 0.5 mM isopropyl b-D-1-thiogalactopyranoside (IPTG) for 12 h at 15°C. The bacteria were harvested by centrifugation, resuspended in Buffer A (50 mM Tris-HCl pH 8.0, 500 mM NaCl, 15 mM imidazole, 0.1% NP-40, 2 mM β-mercaptoethanol and 10% glycerol). 1 mg/ml lysozyme, 5 μg/ml DNase I, 5 mM CaCl_2_ and 25 mM MgCl_2_ (final concentrations) were added to the suspension and the mixture was incubated for 30 min on ice followed by 3 times 20 s sonication at 50% power using the Sonopuls HD 2070 (BANDELIN electronic, Germany, Berlin). The cleared lysate was incubated with Ni-NTA agarose beads (Qiagen, Venlo, Netherlands) for 2 h. The Ni-NTA agarose beads were packed into a plastic column, washed with 10 column volumes of Buffer A and eluted with 2-3 volumes of Buffer A containing 400 mM imidazole. Fractions containing Irc3 fusion protein were pooled and Ulp1 protease was added (10 μg of protease per 1 mg of protein) followed by 12 h dialysis at 4 °C against Buffer B (50 mM Tris-HCl pH 8.0, 500 mM NaCl, 1 mM DTT and 10% glycerol). The dialyzed fractions were diluted to 300 mM NaCl with Buffer O (50 mM Tris-HCl pH 8.0, 1 mM DTT and 10% glycerol), filtered through a 0.22 μm membrane filter and loaded onto a 5 ml HiTrap Heparin HP column (GE Healthcare, Chicago, Illinois, USA). The column was washed with 10 volumes of Buffer H (50 mM Tris-HCl pH 8.0, 300 mM K-Glu, 1 mM DTT and 10% glycerol) and eluted with Buffer F (50 mM Tris-HCl pH 8.0, 1000 mM K-Glu, 1 mM DTT and 10% glycerol). Fractions containing Irc3 were identified by SDS-PAGE, pooled together and stored at −80 °C in small aliquots. The concentration of Irc3 protein was determined spectrophotometrically using a calculated molar extinction coefficient of ε=50770 M^-1^cm^-1^.

### Complementation of the Irc3 disruption phenotype in *Saccharomyces cerevisiae*

A strain of *S. cerevisiae* carrying a deletion of IRC3 [11] was transformed with pRS315-Irc3_op_, pRS315, pRS315-Irc3_sc_ or pRS315-RecG and plated onto glycerol-containing medium to select for the functional mitochondrial genome. The cells were next transferred to a liquid synthetic complete medium without Leu, containing 2% glucose (SC-Leu) for 24, 72 or 168 h. Respiratory-defective and -proficient cells were discriminated by plating onto media containing 0.15% glucose and 3% glycerol.

### ATP hydrolysis activity assay

ATP hydrolysis was measured in 30 mM Tris-HCl pH 8.0, 150 mM K-Glu, 5 mM MgCl_2_, 1 mM DTT, 10% glycerol. The reactions contained 0.1-5 mM ATP, 5-50 nM Irc3 and 5-200 nM a DNA cofactor as indicated and were performed at 37 °C unless specified otherwise (see Suppl. Table S1 for further information on DNA cofactors). The rate of ATP hydrolysis of Irc3 was measured by a conjugated assay linked to NADH oxidation or analyzed from the release of ^32^P orthophosphate using γ-^32^P ATP as a substrate, as previously described [16,19].

### Hydrodynamic characterization of Irc3_op_

The sedimentation velocity of Irc3_op_ was analyzed on a 10-20% glycerol gradient in 30 mM Tris-HCl pH 8.0, 150 mM K-Glu, 1 mM DTT using a MLA-50 rotor in an Optima MAX-XP Ultracentrifuge (Beckman Coulter, Pasadena, California, USA) at 180.000 g for 5 h at 4 °C. The following protein standards of known Svedberg coefficient were used: carbonic anhydrase (2.8S), bovine serum albumin (4.7S) and yeast alcohol dehydrogenase (7.4S). The fractions were analyzed for protein by the Bradford colorimetric assay (Bio-Rad Laboratories, Hercules, California, USA) and for the activity of ATP hydrolysis. Size-exclusion chromatography was performed on a 24 ml Superdex-75 gel filtration column on an ÄKTA Purifier System (GE Healthcare, Chicago, Illinois, USA) using the same buffer as during sedimentation velocity measurement, supplemented with 10% glycerol. The same marker proteins were used for determination of the Stokes radius of Irc3_op_: carbonic anhydrase (*R*_H_ = 2.1 nm), bovine serum albumin (*R*_H_ = 3.6 nm), alcohol dehydrogenase (*R*_H_ = 4.55 nm).

### Analysis of Irc3_op_-DNA complex formation by fluorescence anisotropy

The formation of the Irc3_op_-DNA complex was analyzed in 30 mM Tris-HCl pH 8.0, 150 mM K-Glu, 2 mM MgCl_2_, 1 mM DTT, 1% glycerol, 0.01% NP-40 using 2 nM fluorescein-labeled DNA cofactors. All measurements were performed at 37°C in triplicates. The concentration of Irc3_op_ was varied from 0.5 nM to 1000 nM. Fluorescence anisotropy was measured with a POLARstar Omega plate reader (BMG Labtech, Ortenberg, Baden-Wurttemberg, Germany) in the direct polarization measurement mode and calculated using the following equation:

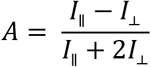

Here, (I_∥_) is the parallel emitted fluorescence signal and (I_⊥_) is the perpendicular emitted fluorescence signal. Data were fitted to a hyperbolic function to calculate the apparent K_d_ using GraphPad 8 software (GraphPad Software, San Diego, California, USA).

### Circular dichroism analysis

Full circular dichroism (CD) spectra analysis as well as the secondary structure melting experiments of Irc3_op_ protein were conducted in 10 mM Potassium Phosphate (pH 7.5), 150 mM (NH_4_)_2_SO_4_ and 1.5 μM Irc3_op_. The spectra were recorded with a Chirascan-plus CD Spectrometer (Applied Photophysics, Leatherhead, Surrey, UK). The CD spectra scanning interval was 180-280 nm. The measured signal was corrected for protein concentration and plotted as molar ellipticity ([θ]). The secondary structure melting experiment was done by continuous recording of the CD signal at 208 nm and gradually increasing temperature by 1 °C/min. Molar ellipticity was plotted as a function of temperature and fitted to an exponential function with a plateau using GraphPad Prism 8 software.

### Helicase unwinding assay of DNA cofactors

The unwinding activity of Irc3_op_ on synthetic DNA cofactors was analyzed as previously described [15]. DNA cofactors were 0.5 nM and the Irc3_op_ protein was 5 nM in all unwinding experiments and the reactions were performed at 37 °C.

## Results

### Irc3 is a mitochondrial helicase conserved in different yeasts

*In silico* analysis by BLAST indicates that the genomes of different yeast species encode a homolog of the Irc3 protein of *Saccharomyces cerevisiae* (Irc3_sc_), previously shown to be a branched-DNA-specific mitochondrial helicase (Fig. 1A, Suppl Fig. 1) [15]. A putative N-terminal mitochondrial targeting peptide that is enriched with hydroxylated and basic amino-acid residues and lacks negatively charged acidic amino-acid residues is present at the N-terminus of Irc3 homologs, suggesting that the members of the Irc3 family are most likely transported to mitochondria. The mitochondrial targeting peptide of Irc3 proteins is followed by the Superfamily II (SF2) helicase domain containing conserved helicase motives (Q, I, Ia, II, III, IV, V, Va, Vb, VI) as defined by Gorbalenya and Koonin [20,21]. The pattern of helicase motifs in is unique in the Irc3-like proteins (Fig. 1B, Suppl. Fig. 1 and Discussion) but the consensus sequences of the motif II and III (DE(H/A)H and SAT, respectively) position Irc3 proteins to SF2 that consists of both RNA and DNA helicases. A distinctive feature of the Irc3 family proteins is a large C-terminal domain that was previously suggested to be involved in the binding of Irc3_sc_ to branched DNA molecules [16]. While five short, conserved elements can be detected in the C-terminal half of Irc3 proteins defined as CI – CV, its structure is less conserved compared to the helicase motor domain, and its size varies from 28 to 36 kDa (Fig. 1B, Suppl. Fig. 1). This led us to the question, how similar are the biochemical properties of Irc3 proteins in different yeast species. We decided to characterize the 68.6 kDa homolog of Irc3_sc_ in the thermotolerant yeast *Ogataea polymorpha* (Irc3_op_) that has a putative mitochondrial targeting signal at the N-terminus followed by a highly conserved helicase domain. However, Irc3_op_ differs substantially from Irc3_sc_ at the C-terminal domain of the protein. A 34-residue deletion between the conserved elements CII and CIII and several shorter stretches of missing residues in the C-terminal half of Irc3_op_ result in a smaller C-terminal domain of Irc3_op_ compared to Irc3_sc_ (29.9 and 36 kDa, respectively Fig. 1A, Suppl Fig. 1). We first asked if Irc3_op_ could complement the *irc3*Δ phenotype in *S. cerevisiae*. Respiratory-proficient yeast cells of *irc3*Δ strains that contain the *wt* mitochondrial genome can be selected for on a media containing a nonfermentable carbon source such as glycerol. However, *irc3*Δ strains rapidly lose their functional mitochondrial genome in the presence of a fermentable carbon source such as glucose, resulting in the formation of respiratory-deficient *petite* cells. To analyze if Irc3_op_ could substitute the endogenous Irc3_sc_, we transformed *W303 irc3*Δ cells with a centromeric plasmid pRS315 expressing either Irc3_sc_, Irc3_op_, or the RecG helicase of *E. coli* and respiratory proficient cells were selected in glycerol containing media. Next, the yeast cultures were released into media containing glucose where the mitochondrial genome of *S. cerevisiae* becomes dispensable (Fig. 1C). The fraction of yeast cells retaining the *wt* mitochondrial DNA was analyzed on media containing low glucose (0.15%) where respiratory active cells containing the *wt* mtDNA form large colonies and respiratory defective petite cells that have lost the functional *wt* mtDNA form small colonies (Fig. 1C). We found that only 5% of *W303 Δirc3* cells transformed with pRS315 were respiratory active after one week of growth in the glucose medium. In contrast, approximately 60% of pRS315-Irc3_op_-transformed cells retained respiratory activity in the same conditions, demonstrating that the expression of plasmid-encoded Irc3_op_ could complement the disruption of the *S. cerevisiae* endogenous IRC3. However, the complementation by Irc3_op_ was not complete as pRS315-Irc3_sc_-transformed cells retained respiratory activity at almost 100% level. In line with previously reported experiments, the bacterial branched DNA-specific RecG helicase of *E. coli* also partially restored the respiratory proficiency of *irc3*Δ yeast cells but less efficiently than Irc3_op_, as only 20% of pRS315-RecG-containing *irc3*Δ cells retained respiratory activity after one week of growth in glucose containing media.

**Figure 1.**
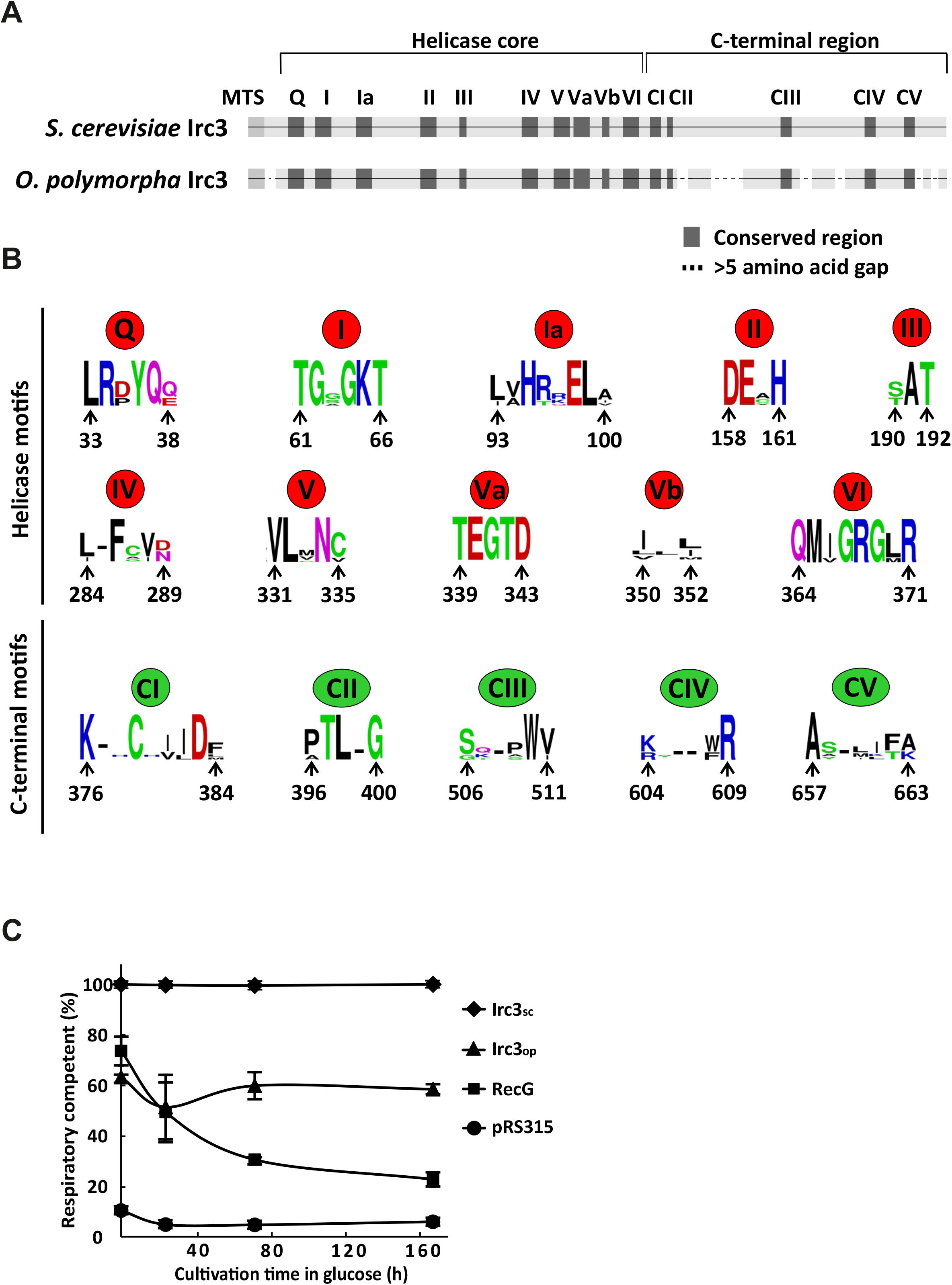
The genome of the thermotolerant yeast *Ogataea polymorpha* encodes a homolog of the *Saccharomyces cerevisiae* helicase Irc3. (A) Schematic alignment of *Saccharomyces cerevisiae* Irc3 (Irc3_sc_) and *Ogataea polymorpha* Irc3 (Irc3_op_). MTS - putative mitochondrial targeting signal. Helicase motives are indicated as Q, I, Ia, II-V, Va, Vb and VI. Irc3-specific C-terminal motives are indicated as CI-CV. A detailed alignment of Irc3 sequences of six yeast species (*Saccharomyces cerevisiae*, *Kluyveromyces lactis*, *Candida albicans*, *Ogataea polymorpha*, *Neurospora crassa* and *Schizosaccharomyces pombe*) is provided in Supplementary Fig. 1. (B) Logos of conserved helicase and C-terminal motives in Irc3 sequence. The letter size corresponds to the level of conservation of the particular amino acid in indicated yeast species. The amino acids are colored according to their side-chains: black – hydrophobic, blue – basic, red – acidic, green – polar, magenta – amids. The numbering of the amino acids corresponds to the sequence of the Irc3_sc_ preprotein. (C) Complementation of the respiratory deficiency phenotype of the *S. cerevisiae Δirc3* strain by Irc3_op_. The fraction of respiratory proficient cells of *S. cerevisiae Δirc3* strain transformed with pRS315, pRS315-Irc3_op_, pRS315-Irc3_sc_, or pRS315-RecG was determined in the course of cultivation in synthetic complete media (-Leu) containing 2% glucose. Each data point is an average of three biological replicas and the error bars indicate the standard deviation of the mean.

### Monomeric Irc3_op_ retains enzymatic activity above 40 °C

To characterize the catalytic properties of Irc3_op_, we expressed the mature mitochondrial form of the helicase in *Escherichia coli* as a His10-Sumo-tagged protein and purified it by a combination of Ni-NTA agarose affinity chromatography, Sumo protease cleavage and heparin agarose chromatography as described in the Materials and Methods (Fig. 2A). The recombinant protein retains full activity for at least 6 months when stored at −70 °C. On a 10-20% glycerol gradient, the recombinant Irc3_op_ sediments as 3.9S (Fig. 2B). The estimated Stokes radius of Irc3_op_ obtained on a Superdex 75 size-exclusion column was 3.85 nm (Fig. 2C). The values of the Irc3_op_ sedimentation coefficient (S) and Stokes radius (R) were used to calculate the apparent molecular weight of Irc3_op_ using the equation:

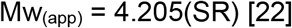

**Figure 2.**
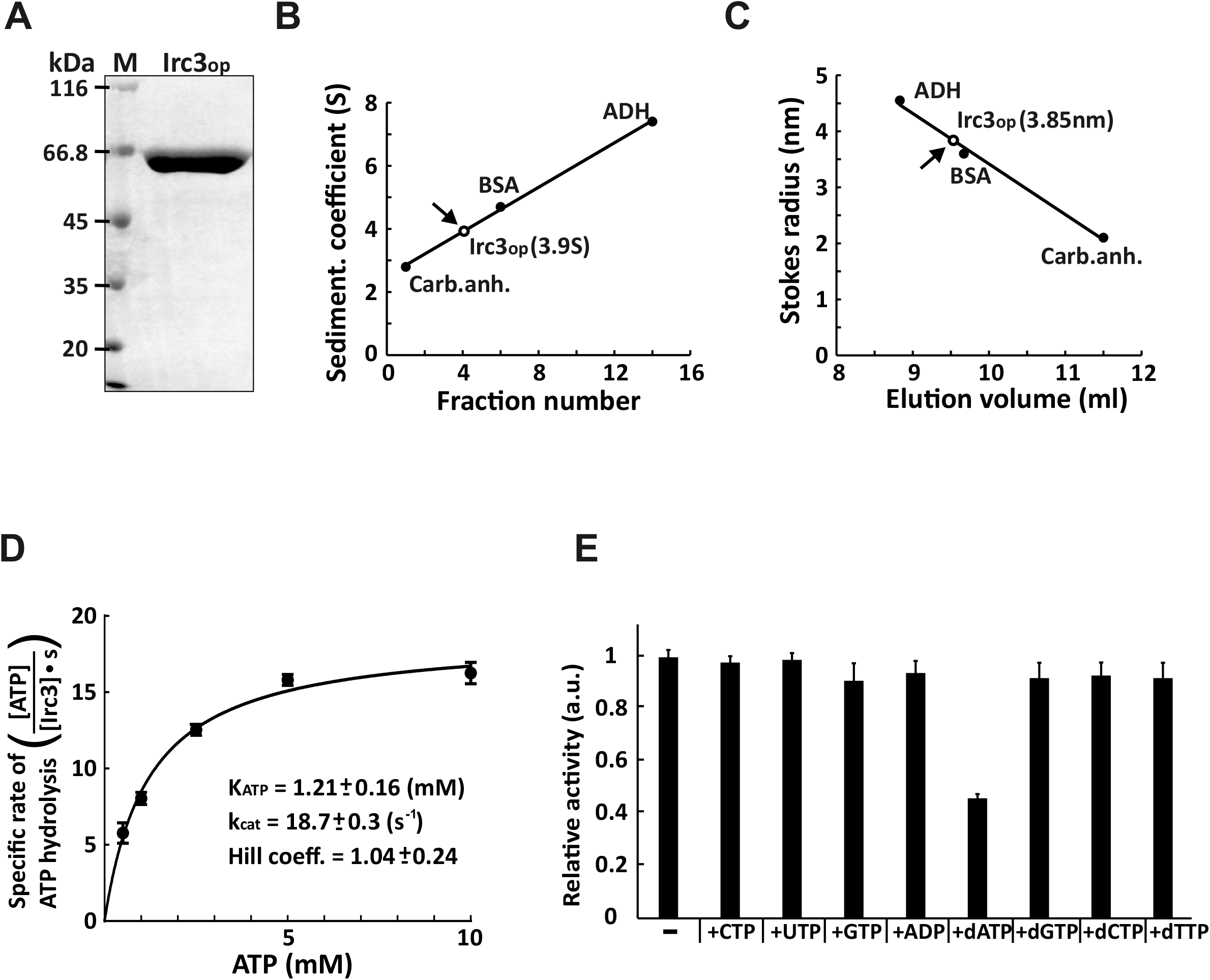
Purification and nucleoside triphosphate hydrolyzing activity of Irc3_op_. (A) Purified Irc3_op_ (15 μg) on a Coomassie stained 10% SDS-PAGE. M – marker proteins with the indicated molecular mass in kilodaltons. (B) Sedimentation of Irc3_op_ (200 μg) and marker proteins (carbonic anhydrase (2.8 S), bovine serum albumin (4.7 S) and alcohol dehydrogenase (7.4 S) on 10 – 20% glycerol gradients. The peak fractions for each protein from the top of the gradient determined by Bradford analysis are indicated. The calculated sedimentation coefficient of Irc3_op_ – 3.9S – is indicated. (C) The mobility of Irc3_op_ and marker proteins carbonic anhydrase (R_H_=2.1 nm), bovine serum albumin (R_H_=3.6 nm) and alcohol dehydrogenase (R_H_=4.55 nm) on a Superdex-75 gel filtration column. The elution volume from sample injection to the protein peak determined by absorbance at 280 nm is shown. The calculated Stokes radius (R) of Irc3_op_ is 3.85 nm. (D) Steady state kinetics of ATP hydrolysis by Irc3 in the presence of 100 nM of a 35 bp dsDNA cofactor. 5 nM Irc3 was titrated by increasing ATP concentrations. The data points representing 3 replicas were fit to Hill equation. The calculated K_M_, k_cat_ and Hill coefficient are indicated. (E) Apparent inhibition of Irc3-catalyzed hydrolysis of 100 μM γ-^32^P-ATP by 5-fold excess of the indicated non-radioactive nucleotide. 100% activity corresponds to a reaction where no additional nucleotides were added. The assays were performed in triplicates and error bars indicate the standard deviation of the mean.

The apparent molecular weight of Irc3_op_ from measured hydrodynamic parameters was 63.1 kDa, which is close to the molecular weight calculated from the amino acid sequence (68.6 kDa). We therefore concluded that Irc3_op_ is a monomeric protein in solution.

Next, basic parameters of ATP hydrolysis catalysis were determined under steady-state conditions at 37 °C in the presence of 100 nM of a 35 bp double-stranded DNA cofactor. Our analysis of ATP hydrolysis reaction kinetics demonstrated that the rate of ATP hydrolysis follows almost perfectly the classical Michaelis-Menten equation (calculated Hill coefficient = 1.04±0.24), demonstrating that Irc3_op_ binds with ATP in a noncooperative manner (Fig. 2D). Based on the assumption that the Irc3_op_ preparation is homogeneous and fully active, the calculated K_M_ value for ATP is 1.21 ±0.16 mM and k_cat_ 18.7±0.3 s^-1^.

We next tested which nucleoside triphosphates can be used as an energy source by Irc3_op_. The hydrolysis rate of γ-^32^P-labelled ATP was analyzed in the presence of a competing nucleoside triphosphate by the charcoal binding assay. We found that Irc3_op_ could utilize efficiently dATP as an energy source (Fig. 2E). However, the other nucleoside triphosphates (CTP, UTP, GTP or dCTP, TTP and dGTP) did not compete significantly with ATP hydrolysis, suggesting that Irc3_op_ preferentially catalyzes the hydrolysis of adenosine nucleoside triphosphates.

*O. polymorpha* is a thermotolerant yeast that can grow at temperatures above 40 °C. We were therefore interested if Irc3_op_ retains its catalytic activity at elevated temperatures and analyzed the kinetics of ATP hydrolysis at different temperatures using a charcoal binding assay. We varied the reaction temperature from 30 to 50 °C in the presence of 100 nM of a 35 bp dsDNA cofactor and found that Irc3_op_ had an optimal temperature for ATP hydrolysis at approximately 41 °C, with more than 75% of enzymatic activity retained at 45 °C (Fig. 3A).

**Figure 3.**
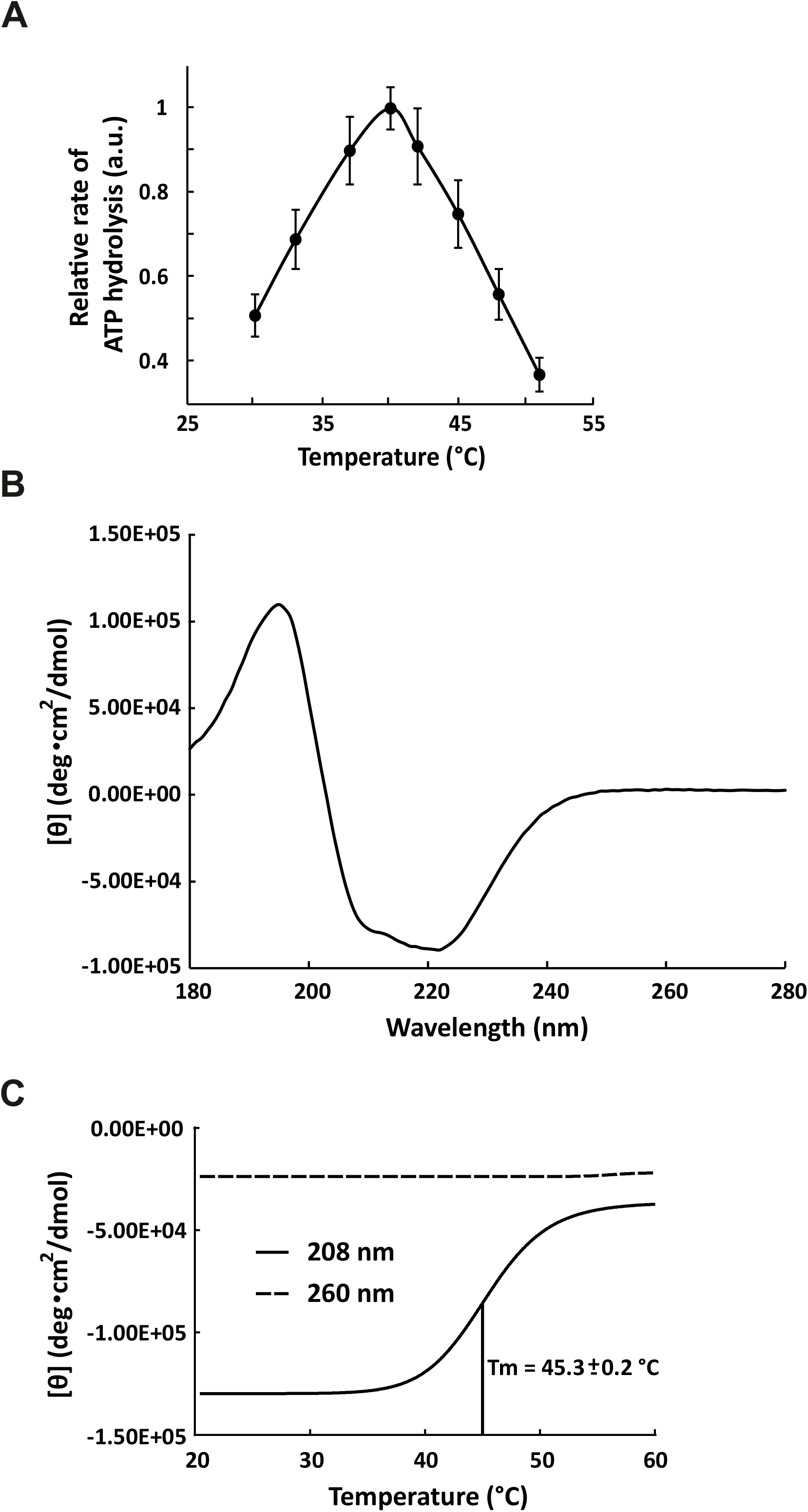
Thermal stability of Irc3_op_. (A) Steady-state rate of ATP hydrolysis as a function of temperature. ATP hydrolysis by Irc3_op_ (50 nM) during 10 min incubation with 100 μM ATP was measured by the release of ^32^P-Pi from γ-^32^P-ATP. Each datapoint is an average of three replicas and the error bars indicate the standard deviation of the mean. (B) CD spectrum of 5 μM Irc3_op_ in the λ=180-280 nm interval measured in 10 mM Potassium Phosphate (pH 7.5), 150 mM (NH_4_)_2_SO_4_. (C) Melting of the secondary structure of Irc3_op_ measured by changes in molar ellipticity at λ=208 nm (indicated by a solid line) during a gradual increase of the temperature by 1 °C/min. Molar ellipticity at λ=260 nm (indicated as a dashed line), used as a control, showed no change in the analyzed range of temperatures.

To further characterize the thermostability of Irc3_op_, we analyzed the CD spectrum of Irc3_op_. The spectrum recorded at room temperature in the λ=180-280 nm interval demonstrates that recombinant Irc3_op_ is properly folded and presumably contains both α and β structures (Fig. 3B). To analyze the stability of the secondary structure, we gradually increased the temperature by 1 °C/min and recorded the CD signal at λ=208 nm that should mostly originate from α-helical structures. The data was fitted to an exponential function using GraphPad Prism 8 software (Fig. 3C). We found that the melting temperature of Irc3_op_ secondary structure is 45.3±0.2 °C. This is in line with the ATP hydrolysis kinetics data that demonstrated that Irc3_op_ retains a significant fraction of its ATPase activity at 45 °C.

### Irc3_op_ binds both single- and double-stranded linear DNA cofactors

We next analyzed Irc3_op_ complex formation with short oligonucleotides. To measure the affinity of Irc3_op_ DNA complex formation, 2 nM of fluorescein-marked DNA was titrated with 1-1000 nM of Irc3_op_ in the absence of ATP (Fig. 4). The titrations that were performed at least in triplicates demonstrated that Irc3_op_ could bind both single-stranded and double-stranded DNA cofactors. The resulting binding data was fitted to a hyperbolic function (Fig. 4A-B) and 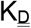 values were deduced, defined as the Irc3 concentrations when 50% of the DNA is in a complex with the Irc3 protein. On Fig. 4C, the K_d_ values are plotted as a function of DNA cofactor length. Our measurements showed that the single-stranded DNA molecules appeared to bind Irc3_op_ slightly more efficiently than double-stranded DNA molecules of the same length. Importantly, all tested single-stranded DNA cofactors with a length of at least 21 nucleotides or double-stranded DNA cofactors that were at least 21 basepairs long bound Irc3_op_ with similar efficiency, characterized by K_d_s of approximately 20 nM and 40 nM, respectively. The 12 nucleotides/basepairs-long linear DNA molecules formed complexes with Irc3 as well, but with lower affinity, suggesting that an optimal binding site for Irc3 consists of 13-20 nucleotides/basepairs.

**Figure 4.**
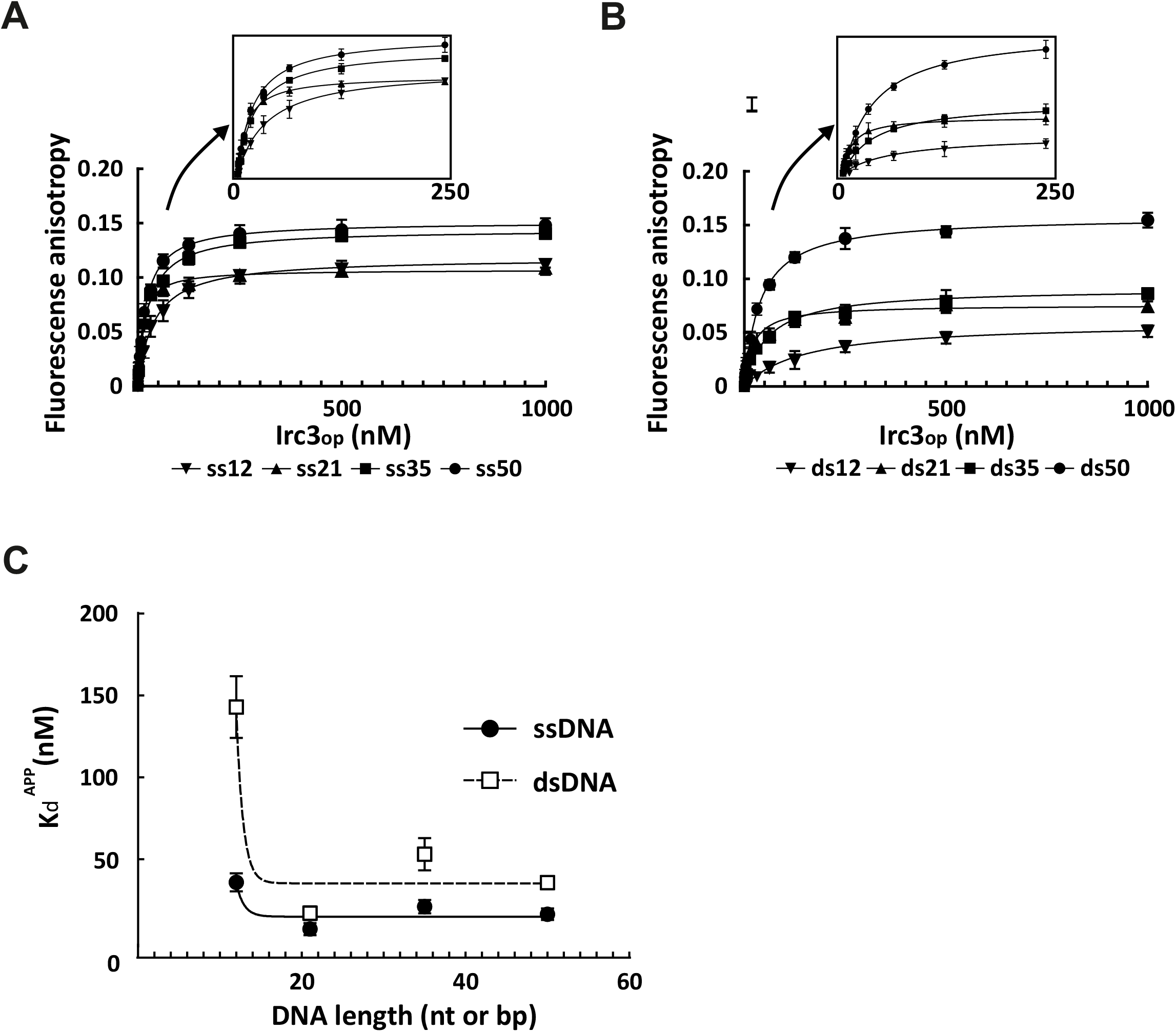
Analysis of Irc3_op_ binding to linear DNA by fluorescence anisotropy. Plot of changes in fluorescence anisotropy upon addition of 0.5 — 1000 nM Irc3_op_ to 2 nM fluorescein-labeled linear DNA. The anisotropy data were fit to a 1:1 binding model. Error bars indicate the standard deviations of three independent replicas. (A) Irc3_op_ binding to single-stranded oligonucleotides of 12 nt (▾), 21 nt (▴), 35 nt (■) and 50 nt (●). (B) Irc3_op_ binding to double-stranded DNA of 12 nt (▾), 21 nt (▴), 35 nt (■) and 50 nt (●). The in-sets on A and B show datapoints expanded between 0 and 250 nM. (C) The data shown in A and B were used to calculate the values of K_d_ for the ssDNA-(●) or dsDNA (□)-Irc3_op_ complexes. K_d_ corresponds to the concentration of total Irc3_op_ when 50% of the DNA is in a complex with the protein. The lines correspond to a best fit of an exponential function with a plateau. Error bars indicate the standard deviation of the mean.

### Irc3_op_ translocates on linear single-stranded DNA and double-stranded DNA

The measurements of Irc3_op_ binding to linear DNA molecules suggested that less than 20 base pairs/nucleotides is required to achieve optimal binding. Even 12mer double- or single-stranded DNA molecules bound Irc3_op_ with a K_d_ that was less than three-fold of that of longer DNA cofactors. To understand if the binding affinity correlates with the stimulation of ATP hydrolysis activity by Irc3_op_, we measured steady-state ATPase kinetics in the presence of single-stranded and double-stranded DNA cofactors ranging from 12 to 75 nucleotides/basepairs using a NADH oxidation conjugated assay. The concentration of Irc3 was held constant at 5 nM, while the molecular concentration of the DNA cofactors was titrated from 5 nM to 200 nM. For each cofactor, the rate of ATP hydrolysis was recorded in triplicates at seven different DNA concentrations, the data was fitted to a hyperbolic function using GraphPad software (Fig. 5A-B) and k_cat_ (Fig. 5C) and K_DNA_ (Fig. 5D) were calculated. k_cat_ corresponds to the rate of ATP hydrolysis obtained at the saturating DNA concentration, divided by the concentration of Irc3. K_DNA_ corresponds to the concentration of a DNA cofactor when the rate of ATP hydrolysis is half of the maximum rate of ATP hydrolysis obtained with the same DNA cofactor. Somewhat surprisingly, the single- and double-stranded DNA cofactors of 12 nucleotides/base pairs showed very little stimulation of ATP hydrolysis by Irc3_op_ in a DNA concentration dependent manner, even though DNA-Irc3_op_ complex formation can be detected at the Irc3_op_ and DNA concentrations used in the assay as indicated by fluorescence polarization measurements (Fig.4C). This suggested that 12 nt/bp is not sufficient for coupling the DNA binding with ATP hydrolysis and presumably not sufficient to start efficient translocation of the helicase on a DNA molecule. All tested DNA cofactors longer than 12 nucleotides/basepairs stimulated ATP hydrolysis by Irc3_op_ in DNA concentration dependent manner. The results showed that K_DNA_ remained roughly constant (Fig. 5C-D), whereas k_cat_ was positively correlated with the length of the DNA cofactor (Fig. 5A-B). According to previously elaborated models [23] the data therefore suggested that Irc3_op_ is a translocating enzyme with the slowest kinetic step related to the enzyme association with DNA, the first isomerization step before processive translocation or the dissociation at the end of the molecule [24].

**Figure 5.**
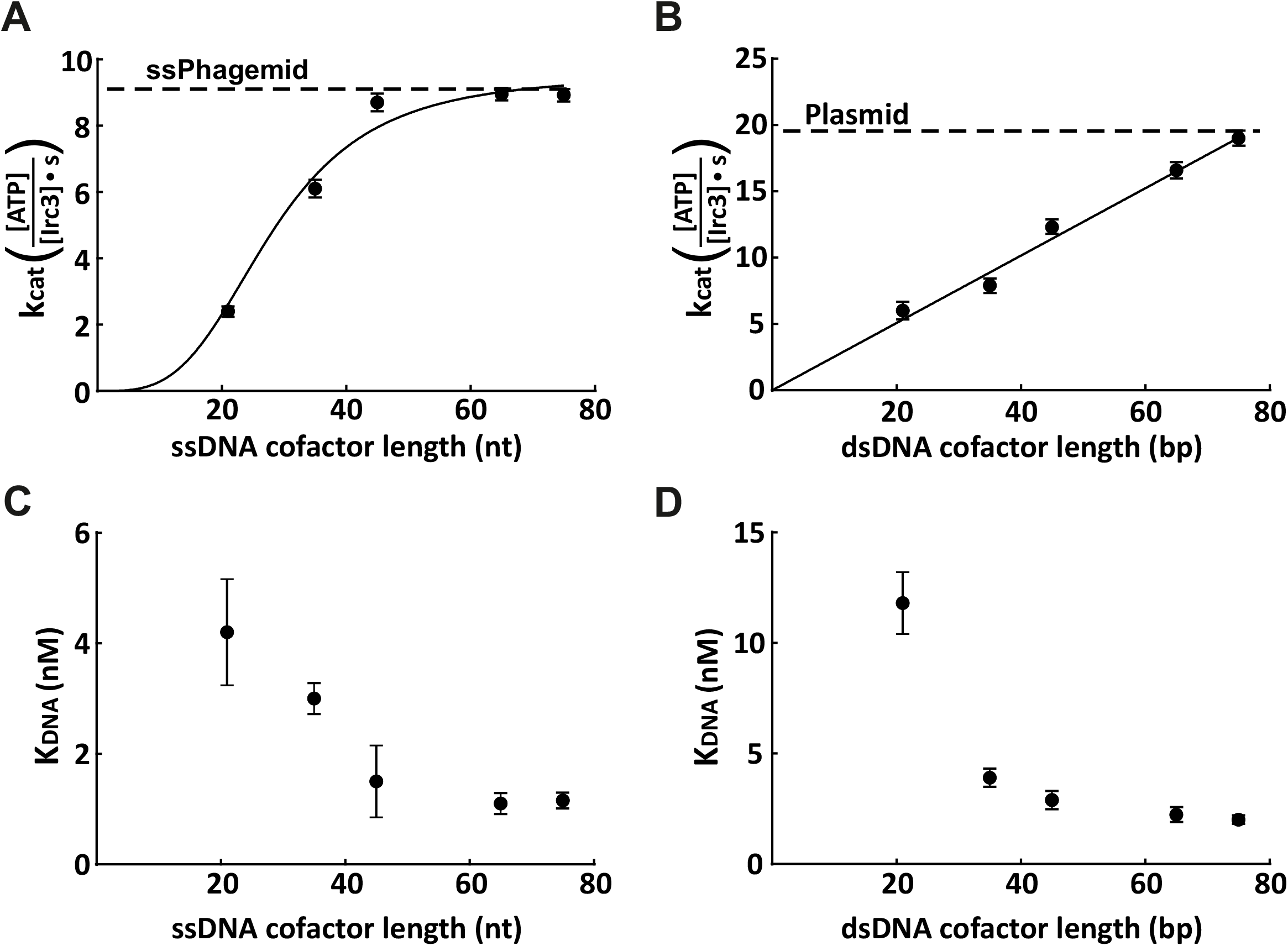
Kinetics of ATP hydrolysis catalyzed by Irc3_op_ in the presence of single-stranded and double-stranded DNA cofactors. Analysis of the steady-state rate of ATP hydrolysis by 5 nM Irc3, measured in the presence of saturating concentrations of (A) ssDNA cofactors of 12, 21, 31, 45, 65 or 75 nt or (B) dsDNA cofactors of 12, 21, 31, 45, 65 or 75 bp. The dashed lines indicate the rate of ATP hydrolysis achieved with the single-stranded phagemid pGEM-11zf+ (A) or with the double-stranded circular plasmid pGEM-11zf+ (B). Analysis of K_DNA_ as a function of a DNA cofactor length. ssDNA cofactors of 12, 21, 31, 45, 65 or 75 nt (C),or dsDNA cofactors of 12, 21, 31, 45, 65 or 75 bp (D). K_DNA_ is the concentration of a DNA cofactor required to achieve a rate of ATP hydrolysis that is 50% of the maximal value with this particular cofactor. Error bars indicate the standard deviation of the mean of three independent replicas.

### Irc3_op_ forms complexes with branched DNA cofactors

Next, we tested the binding of Irc3_op_ to branched DNA cofactors using 2 nM fluorescein-labeled DNA molecules that were titrated with 1-1000 nM Irc3_op_ in the absence of ATP (Fig. 6, Supplementary Fig. S2). The structures of the tested DNA cofactors that were constructed by annealing synthetic oligonucleotides, one of which was carrying a fluorescein moiety at the 5’ end of the molecule, are described in detail in Supplementary Table 2. We analyzed Irc3_op_ binding with three- and four-way branched DNA structures that could mimic possible DNA metabolic intermediates in yeast mitochondria. First, we constructed DNA cofactors representing three-way branched replication forks, including forks with one or both nascent strands missing (Fig 6, columns 1, 2, 5; Supplementary Table 2). Second, we constructed DNA molecules containing specific modifications at the branch-point by introducing a 5 nt gap in the nascent leading or lagging strand (Fig 6, columns 3, 6; Supplementary Table 2) or by the addition of an unpaired 2 nt flap structure (Fig 6, columns 4, 7; Supplementary Table 2). Similar gap- and flap-modifications were introduced into three-way branched cofactors that contained both the nascent leading and lagging strand (Fig. 6, columns 8-16, Supplementary Table 2). Finally, we tested two different four-way junctions: X0, which represents a fixed-structure, and X12, which has a central sliding core (Fig 6, columns 17, 18; Supplementary Table 2).

**Figure 6.**
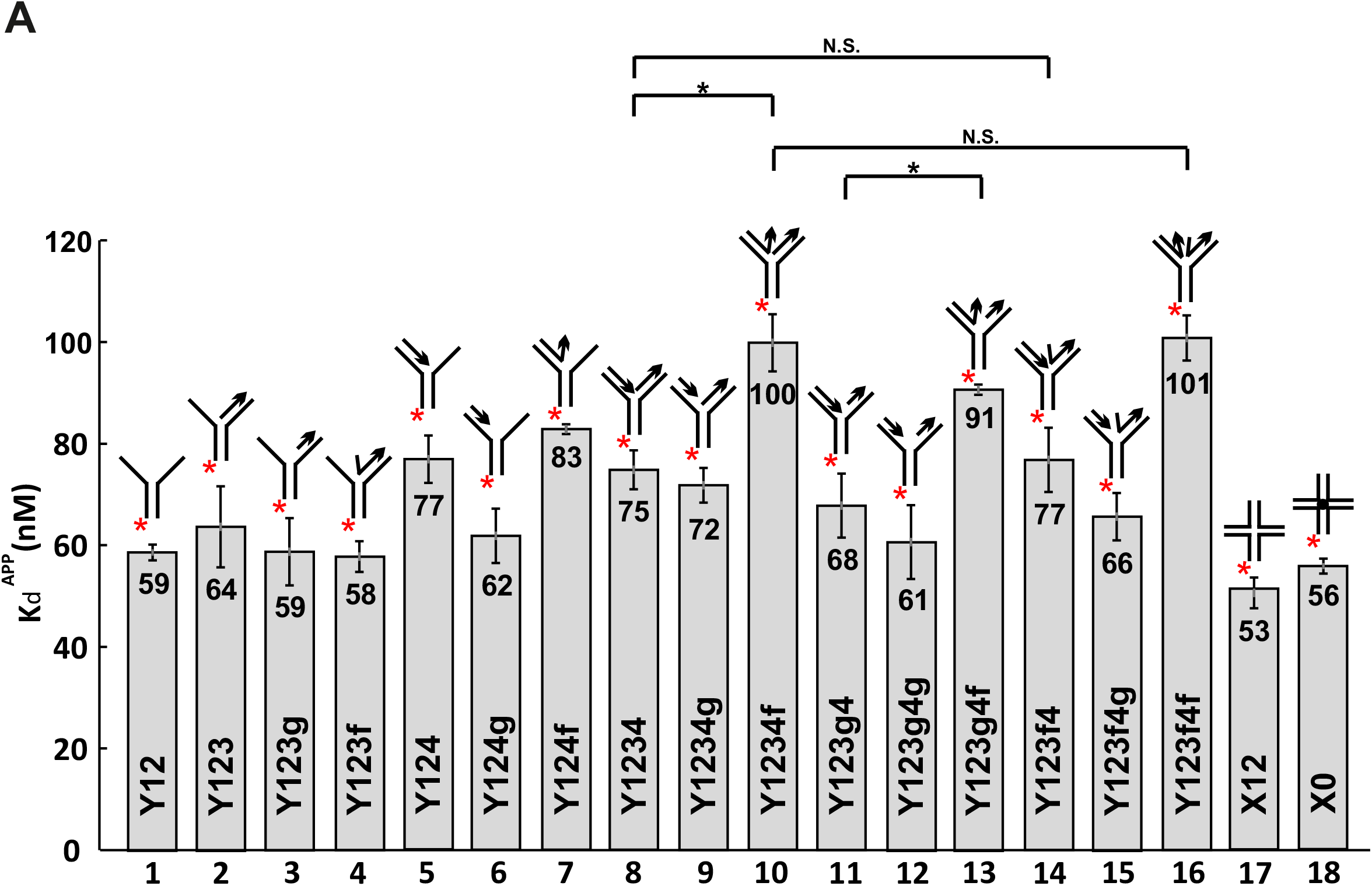
Analysis of Irc3_op_ binding to branched DNA by fluorescence anisotropy. A bar chart showing apparent Kd values for Irc3_op_-DNA complexes mimicking stalled replication forks (the Y cofactors) and a Holliday-junction (the X cofactors), measured by fluorescence anisotropy. Schematic structures of DNA cofactors are depicted above the bars and the cofactors are described in detail in Suppl. Table 2. The names of the Y-series of cofactors indicate the composite oligonucleotides. “g” indicates a 5 nt gap and “h” a 2 nt flap structure. The length of fork branches is 25 nucleotides and the length of the parental double-stranded duplex is 25 bp. The 3’ end of the nascent leading or lagging strand of the fork structure is indicated by an arrow and the position of the fluorescence label is marked with an asterisk. The four-way branched X12 and X0 cofactors contain a mobile or fixed core, respectively, and 25 bp branches [36]. The DNA cofactors are described in detail in Suppl. Table 2. The K_d_ values (nM) for each DNA cofactor measured by fluorescence anisotropy is depicted on each bar. Error bars indicate the standard deviation of the mean of three independent replicas. (*) marks significant difference, p < 0.05, two-tailed Student’s t-test, N.S marks a statistically nonsignificant difference.

Our analysis demonstrated that Irc3_op_ was binding to branched DNA substrates with a K_D_ of approximately 50-100 nM (Fig. 6). Given our previous analysis of Irc3_sc_, demonstrating that different branched DNA substrates are efficiently unwound by the helicase, we expected that Irc3_op_ would bind branched DNA molecules more efficiently than linear DNA molecules. However, we found that the affinity of Irc3_op_ for any branched DNA molecule tested was slightly weaker compared to linear single- or double-stranded DNA molecules that are over 20 nucleotides / base pairs long (Compare Fig. 6 and Fig. 4C). Secondly, we observed a weaker binding of Irc3_op_ to DNA molecules containing a 2-nucleotide flap on the nascent leading strand at the DNA branching point compared to a molecule without a flap This binding difference was detected with branched molecules that contained a complete or gapped lagging strand (Fig. 6, compare columns 8 and 10 or columns 11 and 13). Both the preferential binding of Irc3 to linear DNA molecules compared to branched molecules in general and the inhibition of Irc3 binding by unpaired flap structures suggests that in the absence of ATP, steric contacts of Irc3_op_ with the DNA branch-point interfere with optimal binding.

### Irc3_op_ unwinds branched DNA molecules displaying a preference for replication forks missing the nascent leading strand

We next analyzed the unwinding activity of Irc3_op_ using different model DNA substrates prepared by annealing synthetic oligonucleotides. One of the oligonucleotides used to follow the unwinding reaction was labeled at the 5’-end with ^32^P; detailed structures of the tested DNA structures are described in Supplementary Tables 1 and 2. The model DNA substrates for helicase assays were constructed by annealing the constituent oligonucleotides, which were then purified from polyacrylamide gels and used in unwinding assays at 0.5 nM while the Irc3_op_ helicase was kept at 5 nM, as described in Materials and Methods. The first group of unwinding substrates consists of linear DNA molecules with one strand longer than the other, generating a single-stranded 3’ or 5’ tail (substrates 23 and 14; Fig. 7A, columns 1, 5). The second group consists of substrates that form a three-way branched structure mimicking a replication fork (substrates 123, 123g, 123h, 124, 124g, 124h, 1234; Fig. 7A, columns 2-4 and 6-10). Either the nascent leading- or the nascent lagging strand was labeled in the substrate 1234 (Fig. 7a, columns 9 and 10). The third group consists of substrates that form a four-way junction that mimics a Holliday junction or the product of fork regression (substrates X0, X12; Fig. 7A, columns 11, 12). Both linear DNA substrates tested here in the unwinding assay (substrates 23 and 14) effectively stimulated the ATPase activity of Irc3_op_ (not shown), which was in accordance with experiments analyzing the ATPase activity of Irc3_op_ in the presence of linear single-stranded and double-stranded DNA cofactors (Fig. 5A, B).

**Figure 7.**
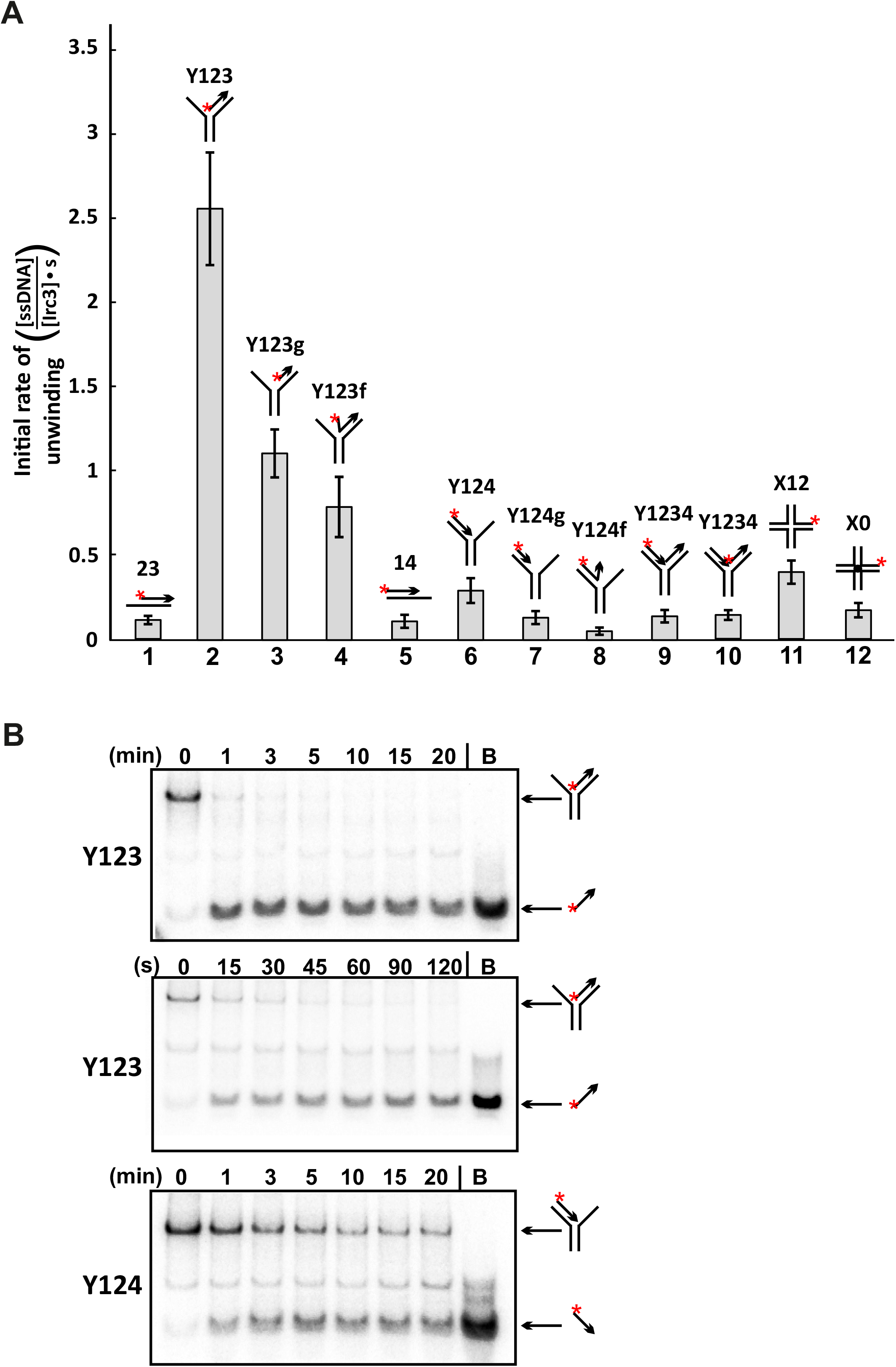
Irc3_op_ strand-separating activity on DNA molecules mimicking a replication fork- and Holliday junctions. (A) A bar chart showing the initial rate of unwinding of different branched DNA substrates. Schematic structures of DNA substrates are depicted above the bars. The 3’ end of the nascent leading or lagging strand of the fork structure is indicated by an arrow and the labelled DNA strand is marked with an asterisk in the name. Substrates marked with a “g” contain a 5 nt gap and substrates marked with an “h” contain a 2 nt unpaired flap. The four-way branched X12 and X0 cofactors contain a mobile or fixed core, respectively, and 25 bp branches [36]. The substrates are described in detail in Suppl. Table 2. Labelled DNA substrates (0.5 nM) and Irc3 (5 nM) were combined and incubated at 37 °C in the presence of Mg^2+^ and ATP for 1, 3, 5, 10, 15 and 20 minutes and for 15, 30, 45, 60, 90 and 120 seconds in case of the substrate 123. The unwinding reactions were stopped by 20 mM EDTA and 0.1% SDS and the products were separated on native polyacrylamide gels and quantified with a phosphorimager. The data were fit to Michaelis Menten kinetics to obtain the value of v_o_. Error bars indicate the standard deviation of the mean of three independent replicas. (B) Images of native polyacrylamide gels used for the calculation of unwinding rates. The two upper panels illustrate the unwinding of a three-way branched substrate mimicking a replication fork with a nascent lagging strand. The lower panel shows the unwinding of a three-way branched substrate mimicking a replication fork with a nascent leading strand. “0” represents a probe incubated for 20 minutes at the reaction temperature in the absence of ATP. “B” – shows the DNA cofactor boiled at 95 °C for 5 min. The time-points at which the samples were removed from the reaction mixture are indicated above the lanes, minutes – in the upper and lower panel; seconds – in the middle panel.

However, our analysis showed that the linear DNA substrates 23 and 14, with a 25 bp double-stranded region and a 25-nt single-stranded overhang tail at the 3’- or 5’-end, were extremely poor substrates for unwinding, resulting in unwinding rates of 0.15 – 0.17 s^-1^ (Fig. 7A, columns 1, 5).

Most DNA substrates of the group 2 mimicking three-way replication forks were substantially more efficient substrates in the helicase reaction compared to the linear DNA molecules (compare columns 1, 2 and 5, 6; Fig. 7A). Within the group 2, a strong preference was observed for forks that contained a nascent lagging strand. For example, substrate 123, which contains a nascent lagging strand, was utilized almost 6-fold faster than substrate 124, which contains a nascent leading strand (compare columns 2 and 6, Fig. 7A and the upper and middle panel with the lower panel on Fig. 7B). Interestingly, the introduction of a 5-nucleotide gap between the branch point and the 5’-end of the nascent lagging strand decreased Irc3_op_ catalytic efficiency almost 2-fold (compare columns 2 and 3, Fig. 7A). Even more prominent was the effect of a 2-nucleotide unpaired flap structure at the 5’-end of the nascent leading strand (compare columns 2 and 4, Fig. 7A). While the catalytic efficiency of Irc3_op_ was significantly lower with DNA substrates that contained the nascent leading strand, a gap or a flap positioned next to the fork branch-point lead to a further diminishment of unwinding activity (compare columns 6 and 7, 8; Fig. 7A). Finally, the catalytic efficiency of Irc3_op_ was low on a fork that contained both the nascent lagging- and leading strand (columns 9 and 10, Fig. 7A).

The substrates of the group 3 mimic four-way Holliday junctions with four double-stranded branches of 25 base pairs. Both the X12 and X0 substrates were remodeled by Irc3_op_, leading to the formation of dimeric DNA molecules (Fig. 7A, columns 11 and 12, and Fig 8A and B). However, the catalytic efficiency of Irc3_op_ on the substrates X12 and X0 was lower than the activity on three-way junctions with a nascent lagging strand (Fig. 7A, columns 11, 12, 2, 3, 4). The four-way junction X12 was utilized slightly more efficiently than X0, which is similar to the preference demonstrated by Irc3_sc_ (Fig. 8A-C). However, while four-way DNA junctions are the best substrates for Irc3_sc_ [15], three-way junctions appeared to be preferred by Irc3_op_ (Fig. 7A, compare columns 2 and 11, 12 and Fig. 8A-C).

**Figure 8.**
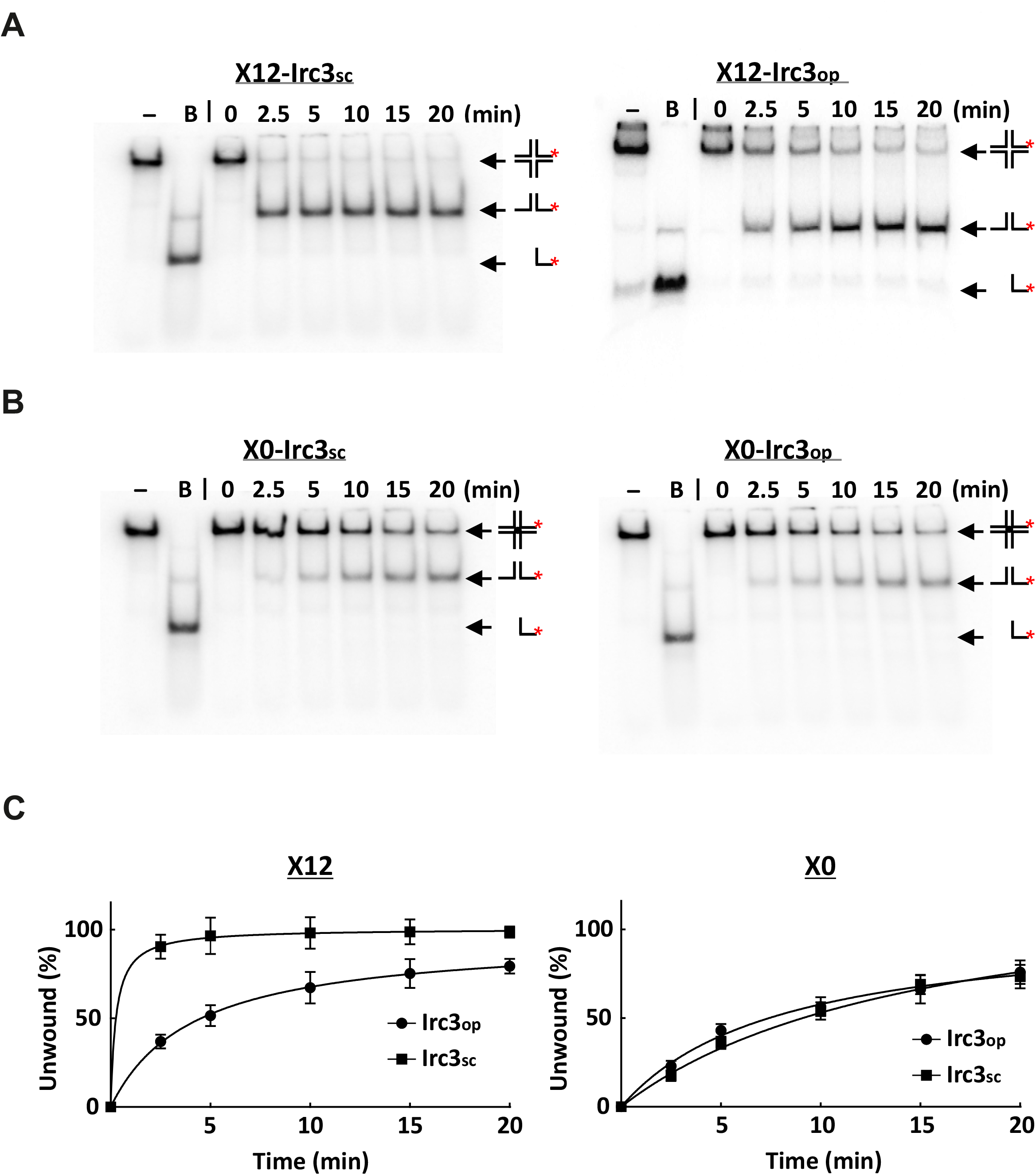
Comparison of unwinding rates of substrates mimicking a Holliday junction with Irc3_sc_ and Irc3_op_. (A and B) Native polyacrylamide gels showing a time-dependent accumulation of unwinding reaction products using Holliday-junction like substrates (0.5 nM) by Irc3_sc_ (5 nM) and Irc3_op_ (5 nM). The mobility of labeled oligonucleotide markers of a monomer, a dimer and a tetramer is indicated with arrows and schematic images on the side of the gels. “–’, Holliday junction substrate, no protein or ATP added, incubated for 20 minutes at the reaction temperature in the absence of ATP. “B” – heat-denatured substrate DNA boiled at 95 °C for 5 min. Different time-points at which the samples were removed from the reaction mixture are indicated above the lanes. Irc3_sc_ (left panels) was incubated at 30 °C and Irc3_op_ (right panels) at 37 °C. A. the X12 substrate; B. X0 substrate. The unwinding reactions were stopped by 20 mM EDTA and 0.1% SDS and the products were separated on native polyacrylamide gels and quantified with a phosphorimager. (C) Quantification of data obtained in A and B presented as graphs showing a time-dependent accumulation of the unwound dimeric product. Unwinding rates using Holliday-junction like substrates X12 (the left graph) and X0 (the right graph) in the reactions catalyzed by Irc3_sc_ (■) and Irc3_op_ (●) are shown. Error bars calculated from three independent experiments show the standard deviations of the mean.

## Discussion

Recent study in our lab has revealed that the Superfamily II helicase Irc3_sc_ is essential for the stability of the mitochondrial genome in the budding yeast *S. cerevisiae* [11]. However, several enzymes involved in mitochondrial DNA metabolism in *S. cerevisiae*, including the helicase Irc3, appear to be relatively thermo-sensitive, which hampers their enzymatic and structural studies. Therefore, we decided to purify a homolog of the *S. cerevisiae* Irc3 from a thermotolerant yeast, as genomic sequencing data indicated that the protein should be present in mitochondria of a variety of yeast species. In this report, we describe the biochemical properties of Irc3_op_, encoded by the open reading frame e_gw1.4.413.1 in *O. polymorpha* (MycoCosm [17]). Our analysis of the purified Irc3_op_ revealed that the Tm of the protein, measured by circular dichroism spectroscopy, is approximately 45 °C, and the optimal temperature for ATP hydrolysis is 41 °C (Fig. 3C). Our unpublished data indicates that Irc3_sc_ is rapidly inactivated at 37 °C, as more than 95% of the ATP hydrolysis activity is lost after 15 minutes of incubation. In general, the helicase domains of Irc3_op_ and Irc3_sc_ are highly similar (approximately 82% over a region of 371 amino acids). However, sequence analysis revealed some distinctive features of Irc3_op_ that could contribute to the better thermotolerance of the protein. There are 13 Pro residues in the helicase domain of Irc3_op_ while only 6 are found in Irc3_sc_. The high number of prolines in Irc3_op_ could contribute to the stiffness of the protein backbone. Furthermore, Irc3_op_ is 6.8 kDa smaller compared to Irc3_sc_ mostly due to deletions in the C-terminal domain and therefore it is possible that flexible loops in the structure of Irc3_op_ are shorter compared to Irc3_sc_ (Fig. 1A). Both the smaller size of the C-terminal domain and the higher content of rigid proline residues in the helicase motor domain could probably contribute to the higher thermotolerance of Irc3_op_ compared to Irc3_sc_.

The Superfamily II helicases show a number of enzymatic activities, including, for example, *bona fide* DNA or RNA helicase activity, nucleic acid translocase activity and chromatin remodeling activity [25,26]. Our analysis indicated that Irc3_op_ is a DNA-dependent ATPase, suggesting that DNA is its natural *in vivo* substrate [11]. To clarify if Irc3_op_ translocates on a DNA molecule or acts locally, we analyzed the dependence of ATP hydrolysis kinetics on the length of DNA cofactor. We found that DNA cofactors stimulate the ATP hydrolysis activity of Irc3_op_ in a length-dependent manner. According to Young et al., 1994, the dependence of k_cat_ and K_DNA_ on the length of nucleic acid cofactors indicates that the DNA-dependent ATPase translocates on the nucleic acid polymer. It is interesting to note that the ATPase activity of Irc3_op_ is stimulated by both dsDNA and ssDNA cofactors suggesting that the helicase could translocate on both dsDNA and ssDNA molecules (Fig. 5A-D). In contrast, our previous experiments suggested that the ATPase activity of Irc3_sc_ is stimulated significantly only by dsDNA cofactors but not by ssDNA [15], suggesting that the interactions between a DNA cofactor and the motor domain of the corresponding helicases might be somewhat different. Our data indicated that the processivity of Irc3_op_ is higher on dsDNA than on ssDNA, as with ssDNA cofactors, the k_cat_ length dependence reached a plateau at 45 nt, but with dsDNA cofactors, the plateau was not reached even with 75 bp (Fig. 5A, B). While not excluding the possibility of a local action mechanism of Irc3_op_ under specific circumstances, these ATP hydrolysis kinetics data strongly suggest that the helicase motor of Irc3_op_ translocates on DNA.

While Irc3_op_ demonstrates the characteristics of a translocating motor enzyme on a linear DNA molecule, the natural unwinding substrate is most likely a DNA molecule that contains a three- or four-way branching point, as demonstrated by the specific unwinding assays with Irc3_op_ described in this report (Fig. 7, 8) and as previously described for Irc3_sc_ [15,16]. Interestingly, Irc3_sc_ and Irc3_op_ proteins display certain differences in their unwinding activity on branched DNA substrates *in vitro*. Irc3_op_ has a clear preference for three-way branched substrates mimicking replication forks that lack the nascent leading strand. On the other hand, Irc3_sc_ is most effective on Holliday junction-like substrates containing a mobile core structure (X12) (Fig. 7 and 8). These findings suggest that the two proteins might have a different substrate preference *in vivo* and that could partially explain why Irc3_op_ is not able to fully compensate for the disruption of the endogenous IRC3 gene in *S. cerevisiae* (Fig. 1C).

The sequence analysis indicates that Irc3 cannot be assigned to any defined helicase family of the Superfamily II, as the combination of conserved helicase motifs in the structure of Irc3 is unique. However, similarly to Irc3, a number of helicases of the Superfamily II resolve different branched DNA molecules and contain additional domains required for the recognition or unwinding of specific branched DNA structures. Remarkably, the members of the RecG and RecQ-like helicase families stand out with their extra domains possessing branched DNA recognition function. Furthermore, it is interesting to note that one member of the RecQ family helicases in human, the RecQL4 helicase, has been suggested to be targeted to mitochondria [27]. RecG of *E. coli*, which can partially complement the loss of Irc3 in yeast mitochondria, has an extra domain called “wedge” at the N-terminus of the protein. The wedge domain has been shown to be important for binding with the branched point of replication fork-like DNA cofactors [28]. Among the RecQ helicase family the C-terminal extra domains such as HRCD (Helicase-and-RNase-D-C-terminal) and RQC (RecQ C-terminal) have been demonstrated to be important for binding to specific substrates [29,30]. Different substrates that have been found to interact with the C-terminal domains of RecQ helicases include, for example, ssDNA for the *E.coli* RecQ helicase and branched DNA for the human WRN helicase [31,32]. However, it is important to note that the C-terminal domain of Irc3 does not show homology to the HRDC and RQC domains in RecQ proteins and its role is largely unexplored.

It remains unclear how Irc3 positions itself on branched DNA and several options can be considered. Irc3 consists of a helicase motor translocating on DNA and our data suggest that the protein makes stronger contacts with the DNA strand that has 3’-5’ polarity in the parental duplex of the forked substrate [16,33]. It is possible that the helicase core domain positions on the parental duplex similarly to what has been suggested for the RecG helicase of *E. coli* [34]. This model explains the formation of a dominant DNA dimeric product during the unwinding of synthetic four-way junction substrates by Irc3_op_ (Fig. 8A, B). However, it can also not be ruled out that similarly to some models describing the action of RecQ family enzymes, the N-terminal helicase motor domain of Irc3 is positioned on one of the DNA branches contacting the nascent lagging strand template. Some members of the RecQ family have been shown to unwind DNA molecules containing a 5’ flap structure, a mechanism that is crucial for strand displacement DNA synthesis in base excision repair or during Okazaki fragment maturation [35]. Our data described here suggest that the flap or gap structure at the DNA branch point inhibits the Irc3 helicase, demonstrating the importance of the specific structure of the DNA junction and implying a difference in Irc3 function compared to RecQ helicases. The alternative mechanisms of Irc3-DNA complex formation proposed here await further functional and structural studies.

## Acknowledgements

We would like to thank Maie Loorits for excellent technical help and Laura Sedman for English Language editing. We thank Dr. Arnold Kristjuhan, Priit Väljamäe and Dr. Jaanus Remme for valuable discussions and help during the study.

## Author Contribution

JS and AF designed the experiments. VP, NG, TS, MR and SS performed the experiments. VP. JS. AF, TS and SS analyzed data. VP and JS wrote the first draft. All authors edited the manuscript.

## SUPPLEMENTAL DATA

### List of Supplementary Materials

1. Supplementary Table S1. List of oligonucleotides used in the study

2. Supplementary Table S2. Model DNA cofactors used in this study

3. Supplementary Figure S1. In silico analysis of Irc3 from different yeast species

4. Supplementary Figure S2. Irc3 complex formation with branched DNA by fluorescence polarization

**Supplementary Table S1.**
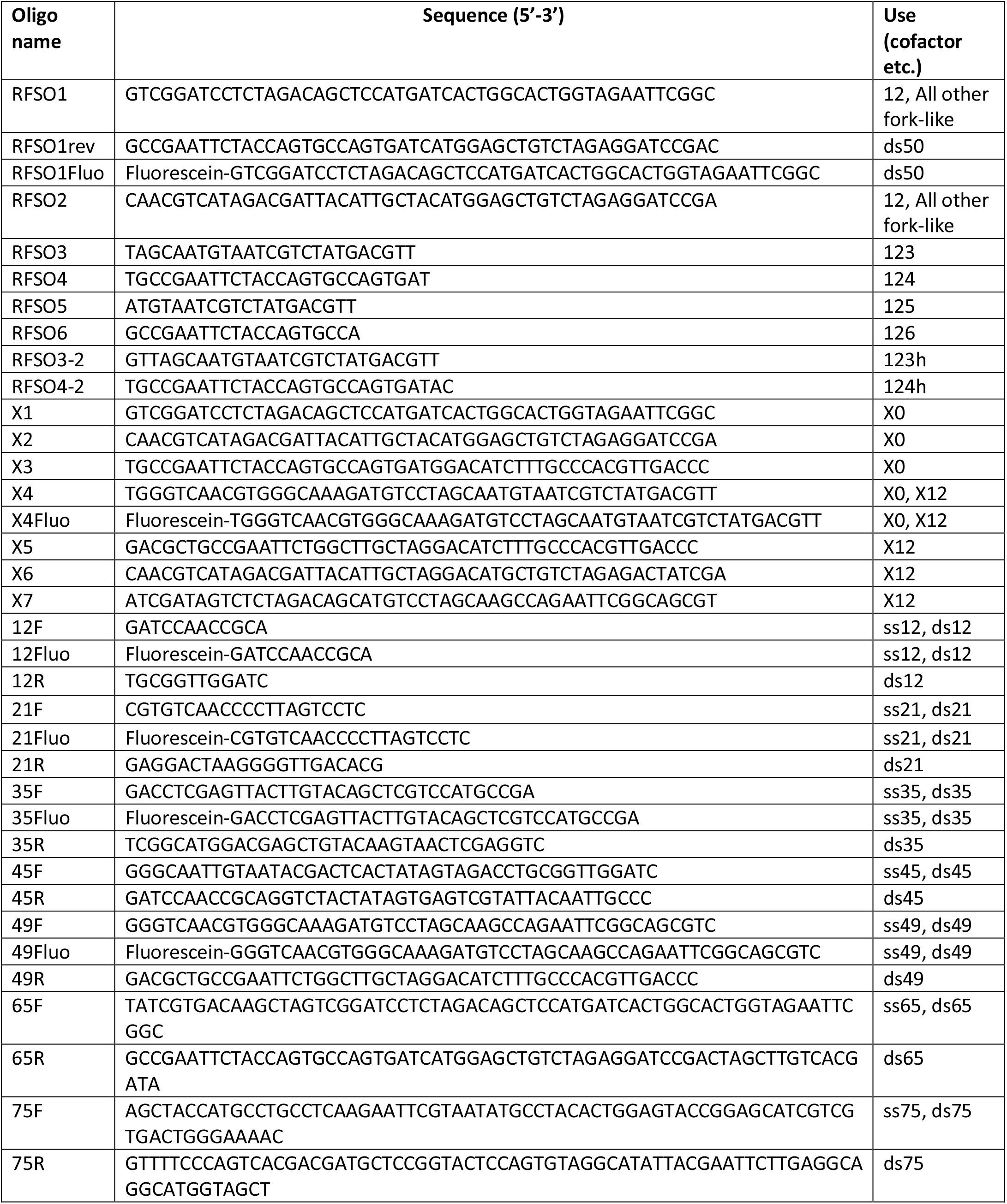
Oligonucleotides used in the study

**Supplementary Table S2.**
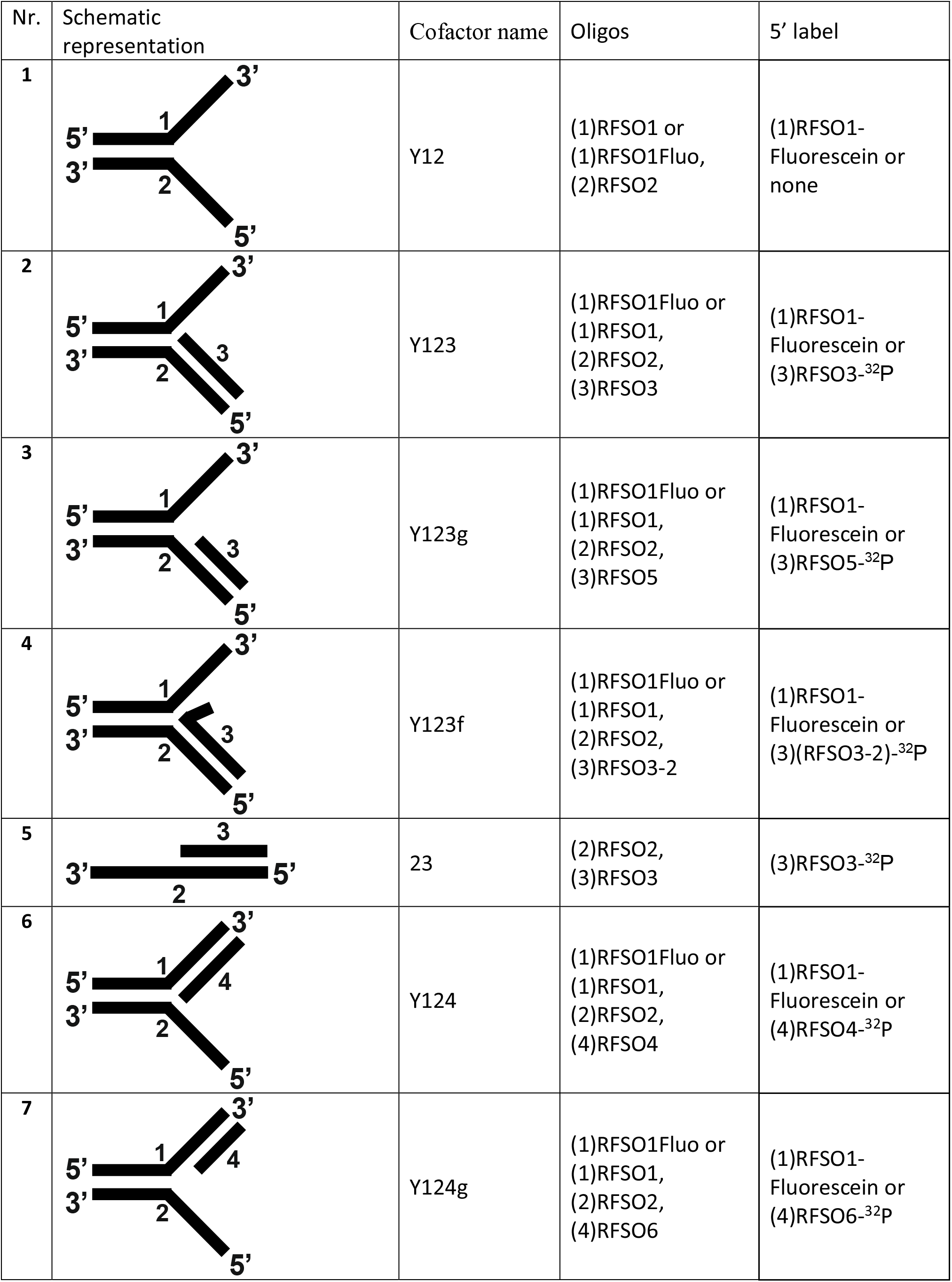
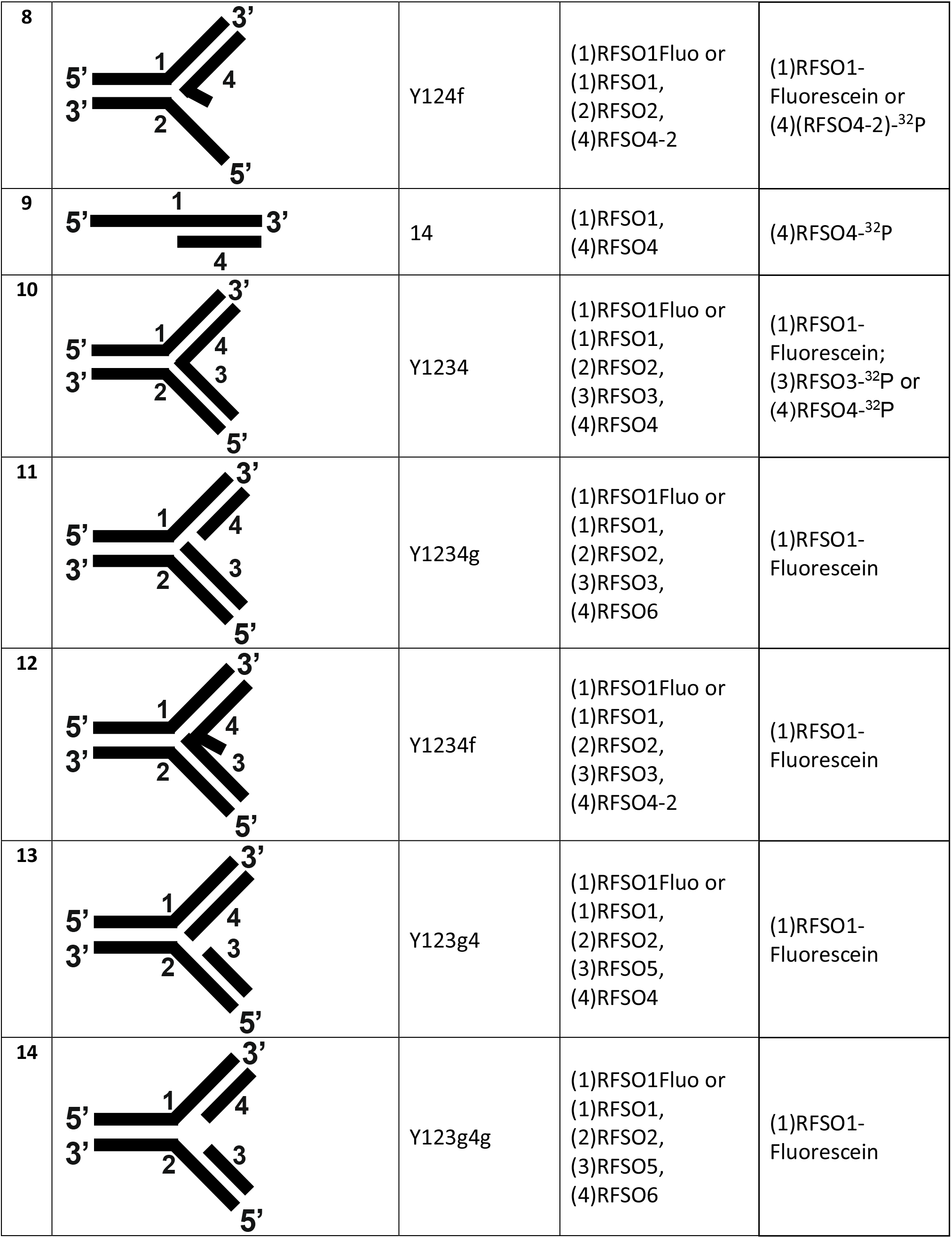
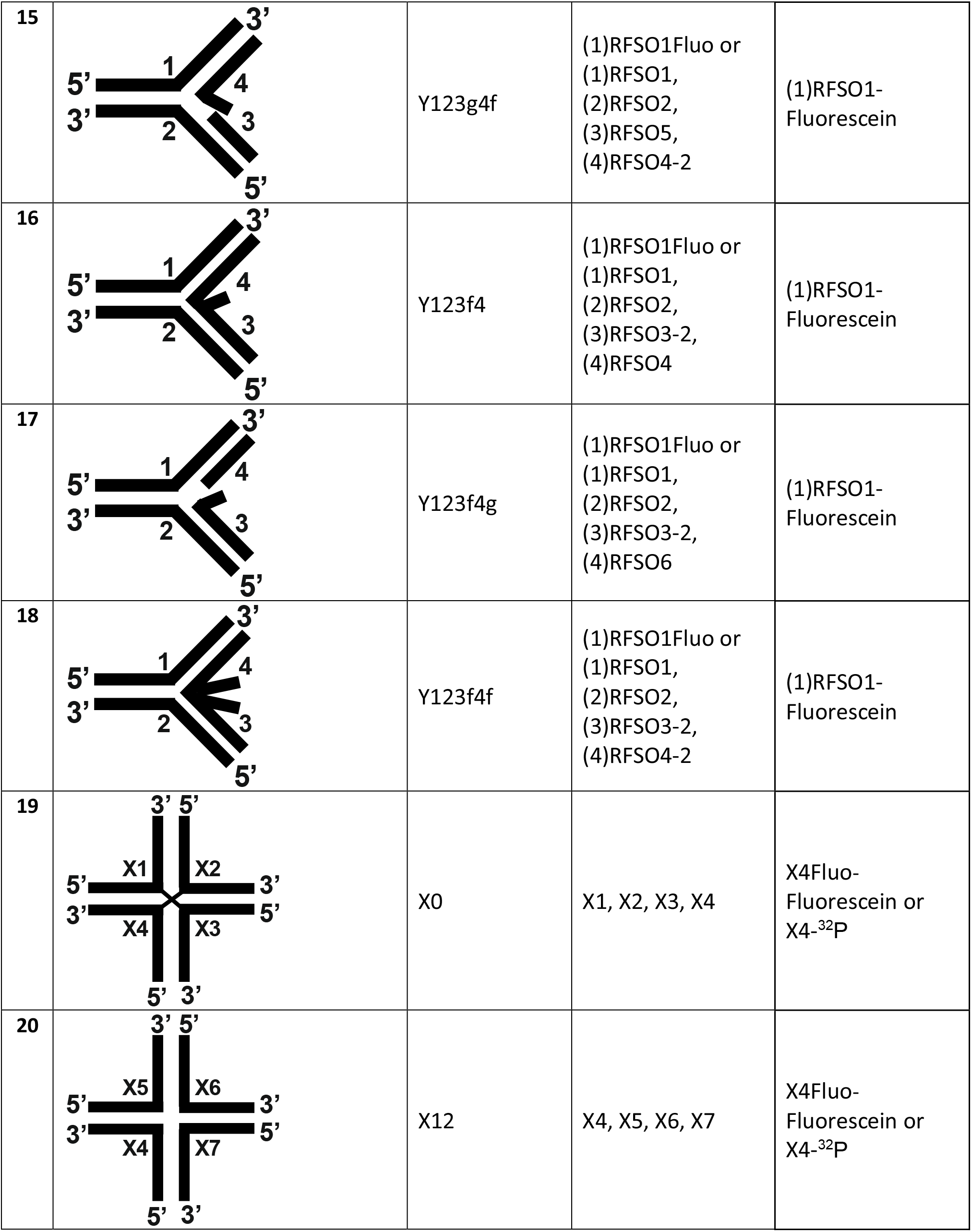
Model DNA cofactors used in this study

Model DNA cofactors used in this study
| Nr. | Schematic representation | Cofactor name | Oligos | 5' label |
| --- | --- | --- | --- | --- |
| 1 |  | Y12 | (1)RFSO1 or<br>(1)RFSO1Fluo,<br>(2)RFSO2 | (1)RFSO1-Fluorescein or<br>none |
| 2 |  | Y123 | (1)RFSO1Fluo or<br>(1)RFSO1,<br>(2)RFSO2,<br>(3)RFSO3 | (1)RFSO1-Fluorescein or<br>(3)RFSO3-<sup>32</sup>P |
| 3 |  | Y123g | (1)RFSO1Fluo or<br>(1)RFSO1,<br>(2)RFSO2,<br>(3)RFSO5 | (1)RFSO1-Fluorescein or<br>(3)RFSO5-<sup>32</sup>P |
| 4 |  | Y123f | (1)RFSO1Fluo or<br>(1)RFSO1,<br>(2)RFSO2,<br>(3)RFSO3-2 | (1)RFSO1-Fluorescein or<br>(3)(RFSO3-2)-<sup>32</sup>P |
| 5 |  | 23 | (2)RFSO2,<br>(3)RFSO3 | (3)RFSO3-<sup>32</sup>P |
| 6 |  | Y124 | (1)RFSO1Fluo or<br>(1)RFSO1,<br>(2)RFSO2,<br>(4)RFSO4 | (1)RFSO1-Fluorescein or<br>(4)RFSO4-<sup>32</sup>P |
| 7 |  | Y124g | (1)RFSO1Fluo or<br>(1)RFSO1,<br>(2)RFSO2,<br>(4)RFSO6 | (1)RFSO1-Fluorescein or<br>(4)RFSO6-<sup>32</sup>P |
| 8 |  | Y124f | (1)RFSO1Fluo or<br>(1)RFSO1,<br>(2)RFSO2,<br>(4)RFSO4-2 | (1)RFSO1-Fluorescein or<br>(4)(RFSO4-2)-<sup>32</sup>P |
| 9 |  | 14 | (1)RFSO1,<br>(4)RFSO4 | (4)RFSO4-<sup>32</sup>P |
| 10 |  | Y1234 | (1)RFSO1Fluo or<br>(1)RFSO1,<br>(2)RFSO2,<br>(3)RFSO3,<br>(4)RFSO4 | (1)RFSO1-Fluorescein;<br>(3)RFSO3-<sup>32</sup>P or<br>(4)RFSO4-<sup>32</sup>P |
| 11 |  | Y1234g | (1)RFSO1Fluo or<br>(1)RFSO1,<br>(2)RFSO2,<br>(3)RFSO3,<br>(4)RFSO6 | (1)RFSO1-Fluorescein |
| 12 |  | Y1234f | (1)RFSO1Fluo or<br>(1)RFSO1,<br>(2)RFSO2,<br>(3)RFSO3,<br>(4)RFSO4-2 | (1)RFSO1-Fluorescein |
| 13 |  | Y123g4 | (1)RFSO1Fluo or<br>(1)RFSO1,<br>(2)RFSO2,<br>(3)RFSO5,<br>(4)RFSO4 | (1)RFSO1-Fluorescein |
| 14 |  | Y123g4g | (1)RFSO1Fluo or<br>(1)RFSO1,<br>(2)RFSO2,<br>(3)RFSO5,<br>(4)RFSO6 | (1)RFSO1-Fluorescein |
| 15 |  | Y123g4f | (1)RFSO1Fluo or<br>(1)RFSO1,<br>(2)RFSO2,<br>(3)RFSO5,<br>(4)RFSO4-2 | (1)RFSO1-Fluorescein |
| 16 |  | Y123f4 | (1)RFSO1Fluo or<br>(1)RFSO1,<br>(2)RFSO2,<br>(3)RFSO3-2,<br>(4)RFSO4 | (1)RFSO1-Fluorescein |
| 17 |  | Y123f4g | (1)RFSO1Fluo or<br>(1)RFSO1,<br>(2)RFSO2,<br>(3)RFSO3-2,<br>(4)RFSO6 | (1)RFSO1-Fluorescein |
| 18 |  | Y123f4f | (1)RFSO1Fluo or<br>(1)RFSO1,<br>(2)RFSO2,<br>(3)RFSO3-2,<br>(4)RFSO4-2 | (1)RFSO1-Fluorescein |
| 19 |  | X0 | X1, X2, X3, X4 | X4Fluo-Fluorescein or<br>X4-<sup>32</sup>P |
| 20 |  | X12 | X4, X5, X6, X7 | X4Fluo-Fluorescein or<br>X4-<sup>32</sup>P |

**Supplementary Figure S1.**
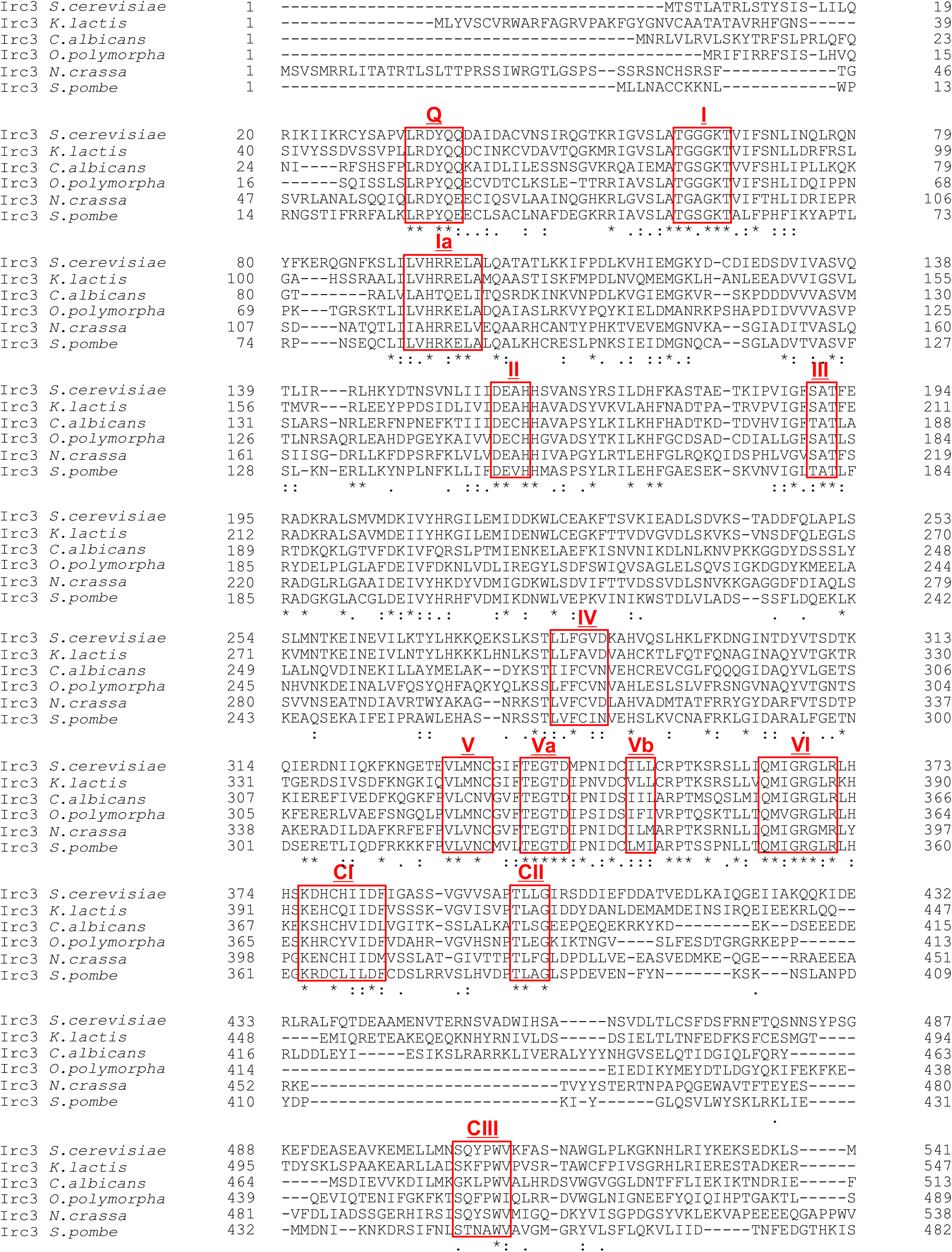
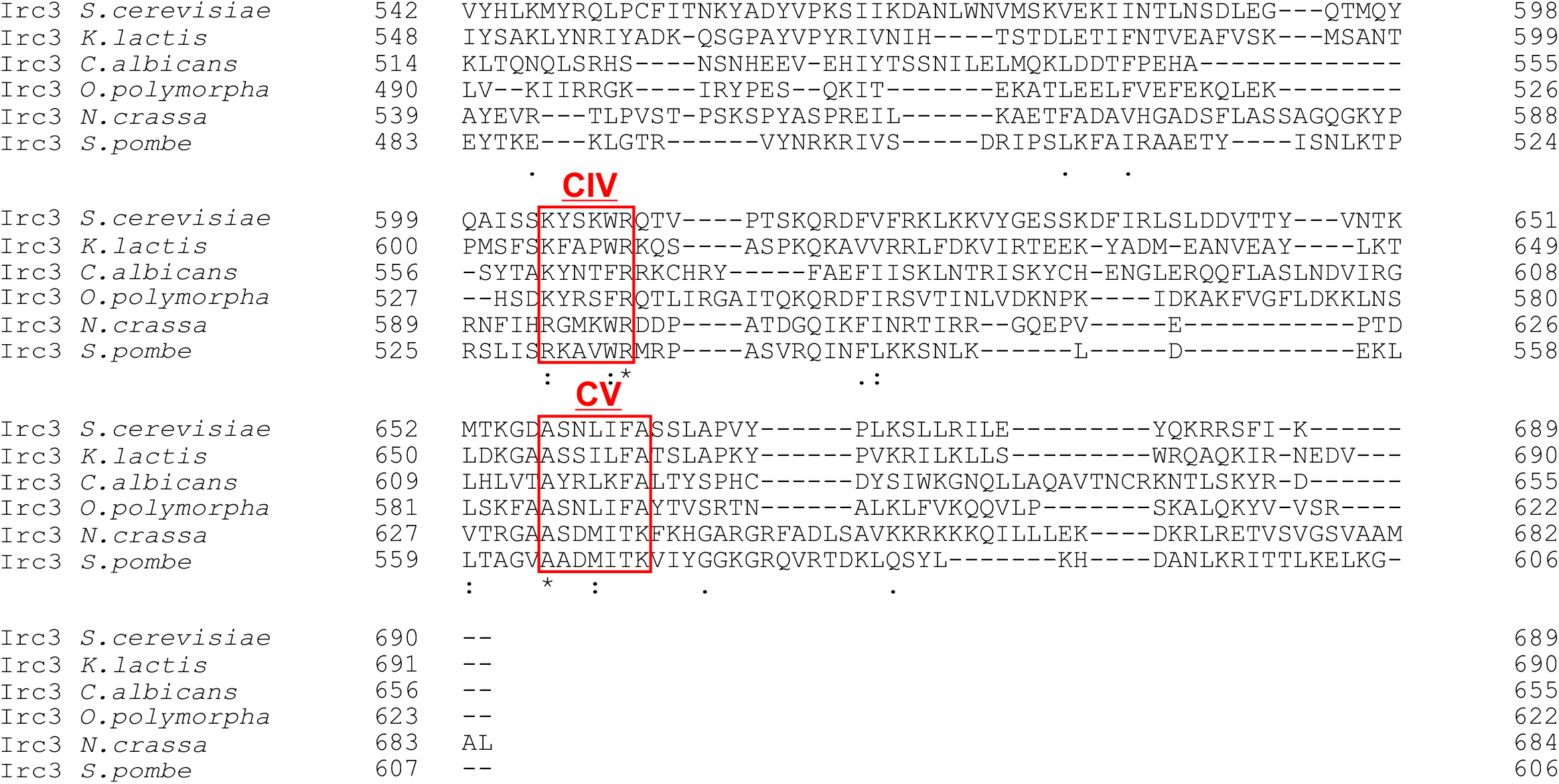
*In silico* analysis of Irc3 from different yeast species. BLAST alignment of Irc3 sequence of different yeast species. Irc3 of the following six yeast species was used in the analysis: *Saccharomyces cerevisiae* (UniProt ref.: Q06683), *Kluyveromyces lactis* (UniProt ref.: Q6CMW9), *Candida albicans* (NCBI ref.: XP_711740.1), *Ogataea polymorpha* (MycoCosm ref.: e_gw1.4.413.1), *Neurospora crassa* (UniProt ref.: Q7RW66), *Schizosaccharomyces pombe* (UniProt ref.: Q1MTR1). “*” (asterisk) indicates positions where all aligned proteins have identical amino acid residue. “:” (colon) indicates conservation of highly similar chemical and physical properties. “.” (period) indicates conservation weakly similar chemical or physical properties.

**Supplementary Figure S2.**
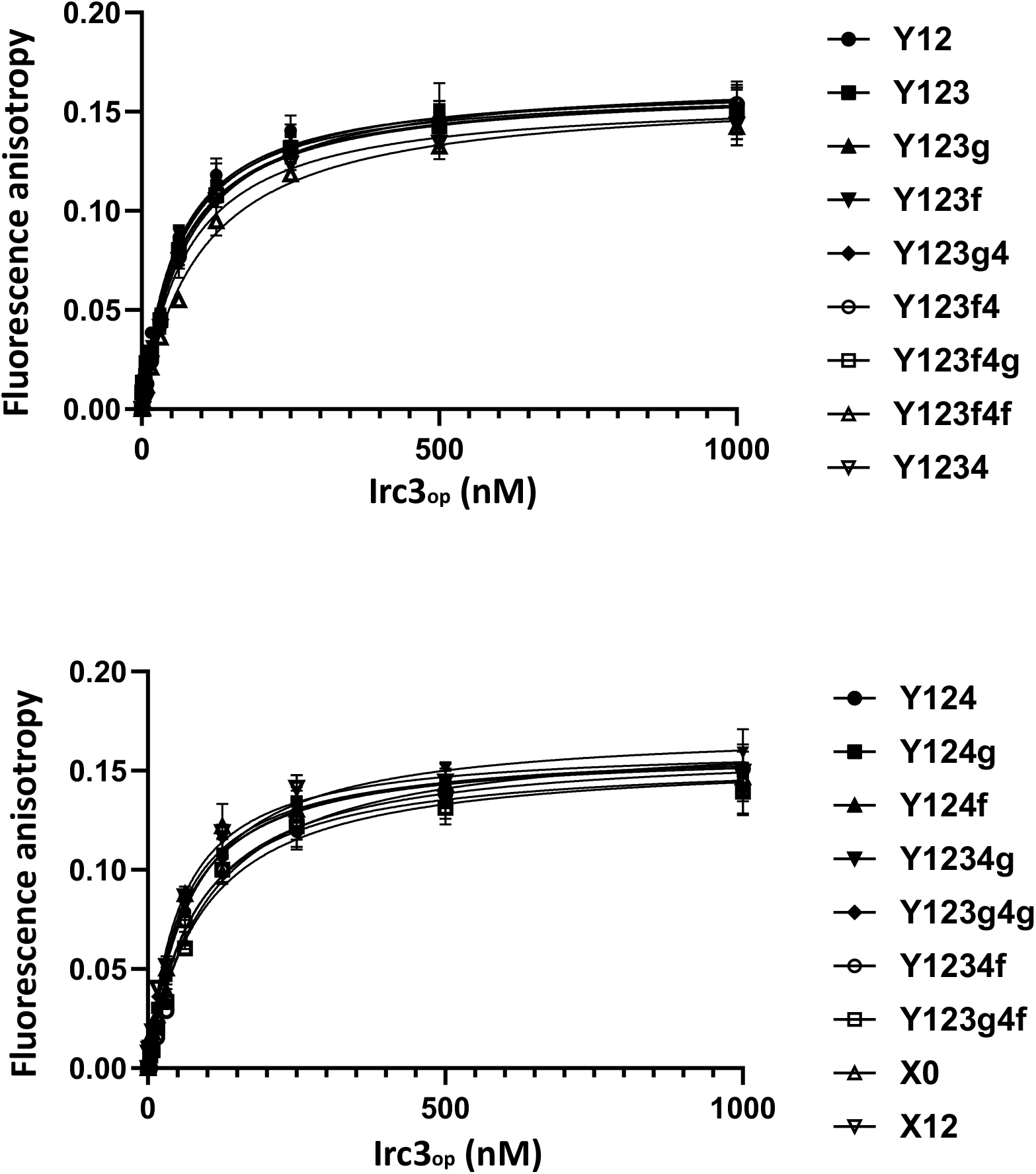
Irc3 complex formation with branched DNA by fluorescence polarization. The plots show changes in fluorescence anisotropy upon addition of 0.5-1000 nM Irc3_op_ to 2 nM fluorescein-labeled branched DNA. The cofactors are described in more detail in Supplementary Table S2. The lines correspond to a best fit of a hyperbolic function. Error bars indicate standard deviations of three independent experiments.

## References

1 Gao Y & Yang W (2020) Different mechanisms for translocation by monomeric and hexameric helicases. Curr Opin Struct Biol 61, 25–32.

2 Singleton MR, Dillingham MS & Wigley DB (2007) Structure and mechanism of helicases and nucleic acid translocases. Annu Rev Biochem 76, 23–50.

3 Ellis NA, Groden J, Ye TZ, Straughen J, Lennon DJ, Ciocci S, Proytcheva M & German J (1995) The Bloom’s syndrome gene product is homologous to RecQ helicases. Cell 83, 655–666.

4 Yu CE, Oshima J, Fu YH, Wijsman EM, Hisama F, Alisch R, Matthews S, Nakura J, Miki T, Ouais S, Martin GM, Mulligan J & Schellenberg GD (1996) Positional cloning of the Werner’s syndrome gene. Science 272, 258–262.

5 Kitao S, Shimamoto A, Goto M, Miller RW, Smithson WA, Lindor NM & Furuichi Y (1999) Mutations in RECQL4 cause a subset of cases of Rothmund-Thomson syndrome. Nat Genet 22, 82–84.

6 Su Y, Meador JA, Calaf GM, DeSantis LP, Zhao Y, Bohr VA & Balajee AS (2010) Human RecQL4 helicase plays critical roles in prostate carcinogenesis. Cancer Res 70, 9207–9217.

7 Arai A, Chano T, Futami K, Furuichi Y, Ikebuchi K, Inui T, Tameno H, Ochi Y, Shimada T, Hisa Y & Okabe H (2011) RECQL1 and WRN Proteins Are Potential Therapeutic Targets in Head and Neck Squamous Cell Carcinoma. Cancer Res 71, 4598–4607.

8 Song J, Herrmann JM & Becker T (2021) Quality control of the mitochondrial proteome. Nat Rev Mol Cell Biol 22, 54–70.

9 Lahaye A, Stahl H, Thines-Sempoux D & Foury F (1991) PIF1: a DNA helicase in yeast mitochondria. EMBO J 10, 997–1007.

10 Sedman T, Kuusk S, Kivi S & Sedman J (2000) A DNA Helicase Required for Maintenance of the Functional Mitochondrial Genome in Saccharomyces cerevisiae. Mol Cell Biol 20, 1816–1824.

11 Sedman T, Gaidutšik I, Villemson K, Hou Y & Sedman J (2014) Double-stranded DNA-dependent ATPase Irc3p is directly involved in mitochondrial genome maintenance. Nucleic Acids Res 42, 13214–13227.

12 Zhang D-H, Zhou B, Huang Y, Xu L-X & Zhou J-Q (2006) The human Pif1 helicase, a potential Escherichia coli RecD homologue, inhibits telomerase activity. Nucleic Acids Res 34, 1393–1404.

13 Futami K, Shimamoto A & Furuichi Y (2007) Mitochondrial and nuclear localization of human Pif1 helicase. Biol Pharm Bull 30, 1685–1692.

14 Spelbrink JN, Li FY, Tiranti V, Nikali K, Yuan QP, Tariq M, Wanrooij S, Garrido N, Comi G, Morandi L, Santoro L, Toscano A, Fabrizi GM, Somer H, Croxen R, Beeson D, Poulton J, Suomalainen A, Jacobs HT, Zeviani M & Larsson C (2001) Human mitochondrial DNA deletions associated with mutations in the gene encoding Twinkle, a phage T7 gene 4-like protein localized in mitochondria. Nat Genet 28, 223–231.

15 Gaidutšik I, Sedman T, Sillamaa S & Sedman J (2016) Irc3 is a mitochondrial DNA branch migration enzyme. Sci Rep 6, 26414.

16 Sedman T, Garber N, Gaidutšik I, Sillamaa S, Paats J, Piljukov VJ & Sedman J (2017) Mitochondrial helicase Irc3 translocates along double-stranded DNA. FEBS Lett 591, 3831–3841.

17 Grigoriev IV, Nikitin R, Haridas S, Kuo A, Ohm R, Otillar R, Riley R, Salamov A, Zhao X, Korzeniewski F, Smirnova T, Nordberg H, Dubchak I & Shabalov I (2014) MycoCosm portal: gearing up for 1000 fungal genomes. Nucleic Acids Res 42, D699–D704.

18 Turnbull KJ, Dzhygyr I, Lindemose S, Hauryliuk V & Roghanian M (2019) Intramolecular Interactions Dominate the Autoregulation of Escherichia coli Stringent Factor RelA. Front Microbiol 10.

19 Kuusk S, Sedman T, Jõers P & Sedman J (2005) Hmi1p from Saccharomyces cerevisiae mitochondria is a structure-specific DNA helicase. J Biol Chem 280, 24322–24329.

20 Gorbalenya AE & Koonin EV (1993) Helicases: amino acid sequence comparisons and structure-function relationships. Curr Opin Struct Biol 3, 419–429.

21 Fairman-Williams ME, Guenther U-P & Jankowsky E (2010) SF1 and SF2 helicases: family matters. Curr Opin Struct Biol 20, 313–324.

22 Erickson HP (2009) Size and shape of protein molecules at the nanometer level determined by sedimentation, gel filtration, and electron microscopy. Biol Proced Online 11, 32–51.

23 Young MC, Kuhl SB & von Hippel PH (1994) Kinetic theory of ATP-driven translocases on one-dimensional polymer lattices. J Mol Biol 235, 1436–1446.

24 Fischer CJ, Saha A & Cairns BR (2007) Kinetic Model for the ATP-Dependent Translocation of Saccharomyces cerevisiae RSC along Double-Stranded DNA. Biochemistry 46, 12416–12426.

25 Pyle AM (2008) Translocation and Unwinding Mechanisms of RNA and DNA Helicases. Annu Rev Biophys 37, 317–336.

26 Thomä NH, Czyzewski BK, Alexeev AA, Mazin AV, Kowalczykowski SC & Pavletich NP (2005) Structure of the SWI2/SNF2 chromatin-remodeling domain of eukaryotic Rad54. Nat Struct Mol Biol 12, 350–356.

27 Croteau DL, Rossi ML, Canugovi C, Tian J, Sykora P, Ramamoorthy M, Wang ZM, Singh DK, Akbari M, Kasiviswanathan R, Copeland WC & Bohr VA (2012) RECQL4 localizes to mitochondria and preserves mitochondrial DNA integrity. Aging Cell 11, 456–466.

28 Briggs GS, Mahdi AA, Wen Q & Lloyd RG (2005) DNA binding by the substrate specificity (wedge) domain of RecG helicase suggests a role in processivity. J Biol Chem 280, 13921–13927.

29 Harami GM, Nagy NT, Martina M, Neuman KC & Kovács M (2015) The HRDC domain of E. coli RecQ helicase controls single-stranded DNA translocation and double-stranded DNA unwinding rates without affecting mechanoenzymatic coupling. Sci Rep 5, 11091.

30 Tadokoro T, Kulikowicz T, Dawut L, Croteau DL & Bohr VA (2012) DNA binding residues in the RQC domain of Werner protein are critical for its catalytic activities. Aging 4, 417–429.

31 von Kobbe C, Thomä NH, Czyzewski BK, Pavletich NP & Bohr VA (2003) Werner syndrome protein contains three structure-specific DNA binding domains. J Biol Chem 278, 52997–53006.

32 Bernstein DA & Keck JL (2005) Conferring substrate specificity to DNA helicases: role of the RecQ HRDC domain. Struct Lond Engl 1993 13, 1173–1182.

33 Piljukov V-J, Garber N, Sedman T & Sedman J (2020) Irc3 is a monomeric DNA branch point-binding helicase in mitochondria of the yeast Saccharomyces cerevisiae. FEBS Lett 594, 3142–3155.

34 McGlynn P & Lloyd RG (2001) Rescue of stalled replication forks by RecG: Simultaneous translocation on the leading and lagging strand templates supports an active DNA unwinding model of fork reversal and Holliday junction formation. Proc Natl Acad Sci U S A 98, 8227–8234.

35 Sharma S, Sommers JA, Choudhary S, Faulkner JK, Cui S, Andreoli L, Muzzolini L, Vindigni A & Brosh RM (2005) Biochemical analysis of the DNA unwinding and strand annealing activities catalyzed by human RECQ1. J Biol Chem 280, 28072–28084.

36 Lloyd RG & Sharples GJ (1993) Dissociation of synthetic Holliday junctions by E. coli RecG protein. EMBO J 12, 17–22.

